# Chloroplast chaperonin-mediated targeting of a thylakoid membrane protein

**DOI:** 10.1101/2020.04.20.051433

**Authors:** Laura Klasek, Kentaro Inoue, Steven M. Theg

## Abstract

Post-translational protein targeting requires chaperone assistance to direct insertion-competent proteins to integration pathways. Chloroplasts integrate nearly all thylakoid transmembrane proteins post-translationally, but mechanisms in the stroma that assist their insertion remain largely undefined. Here, we investigated how the chloroplast chaperonin (Cpn60) facilitated the thylakoid integration of Plastidic type I signal peptidase 1 (Plsp1) using in vitro targeting assays. Cpn60 bound Plsp1 in the stroma. In isolated chloroplasts, the membrane integration of imported Plsp1 correlated with its dissociation from Cpn60. When the Plsp1 residues that interacted with Cpn60 were removed, Plsp1 did not integrate into the membrane. These results suggested Cpn60 was an intermediate in Plsp1’s thylakoid targeting. In isolated thylakoids, the integration of Plsp1 decreased if Cpn60 was present in excess of cpSecA1, the stromal motor of the cpSec1 translocon which inserts unfolded Plsp1 into the thylakoid. An excess of cpSecA1 favored integration. Introducing Cpn60’s obligate substrate RbcL displaced Cpn60-bound Plsp1; then, the released Plsp1 exhibited increased accessibility to cpSec1. These in vitro targeting experiments support a model in which Cpn60 captures and then releases insertion-competent Plsp1, while cpSecA1 recognizes free Plsp1 for integration. Thylakoid transmembrane proteins transiting the stroma can interact with Cpn60 to shield from the aqueous environment.

**One-sentence summary:** The chloroplast chaperonin captures and releases Plastidic type I signal peptidase 1 during its targeting to the thylakoid membrane.

## Introduction

In oxygenic photosynthesis, the light-harvesting reactions and ATP synthesis occur at the thylakoid, the internal membrane of chloroplasts and cyanobacteria. Thylakoid development requires coordinated synthesis, delivery, and assembly of its proteins, pigments, and lipids (Rast et al., 2015). Co-translational integration is the predominant mechanism for inserting membrane proteins into a lipid bilayer when these translation and integration may be spatially coupled. This process targets an estimated 86% and 75% of membrane proteins to the *E. coli* plasma membrane and yeast ER, respectively (Alamo et al., 2011; Schibich et al., 2016). Ribosomes decorate thylakoids in cyanobacteria (Rast et al., 2019), suggesting co-translational integration of cyanobacterial thylakoid proteins. However, in plants, the nuclear genome encodes the majority of integral thylakoid transmembrane (TM) proteins, and the plastid genome encodes about 40 TM proteins. Of approximately 170 thylakoid proteins with at least one predicted TM domain (TMD), only 10% integrate co-translationally (Sun et al., 2009; Zoschke and Barkan, 2015). Chloroplasts depend upon post-translational mechanisms to deliver thylakoid TM proteins.

To facilitate its post-translational targeting, a thylakoid TM protein may need to be folded or kept unfolded in the stroma. The chloroplast secretory (cpSec1) and signal recognition particle (cpSRP) pathways both insert unfolded proteins into the thylakoid (Klasek and Inoue, 2016). During in vitro targeting assays, premature folding of the cpSec1 substrate Plastidic type I signal peptidase 1 (Plsp1) mistargets Plsp1 to the incorrect side of the thylakoid (Endow et al., 2015). The cpSRP substrate light-harvesting complex protein (LHCP) rapidly aggregates (Payan and Cline, 1991; Jaru-Ampornpan et al., 2010). The chloroplast must chaperone unfolded thylakoid TM proteins transiting the stroma.

An unfolded, insertion-competent state can be stabilized by either specific targeting factors or promiscuous chaperones. In *E. coli*, the SecB targeting factor keeps secreted proteins unfolded (Sala et al., 2014). The cytosolic GET pathway targets tail-anchored (TA) proteins to the ER through a cascade of chaperones which increase in specificity for insertion-competent TA substrates (Shan, 2019). In contrast, the mechanisms which stabilize thylakoid proteins in transit are largely unknown beyond two specific examples. First, the plastid-specific cpSRP43 targeting factor assists LHCP in transiting the stroma by preventing and reversing its aggregation (Payan and Cline, 1991; Jaru-Ampornpan et al., 2010). Second, the chloroplast Hsp90 chaperone promotes the thylakoid association of cpSec1-targeted OE33/PsbO (Jiang et al., 2017). In addition, the stromal chaperones Cpn60, ClpC/Hsp93, Hsp70, and Hsp90 all interact with newly imported proteins (Lubben et al., 1989; Madueno et al., 1993; Tsugeki and Nishimura, 1993; Bonk et al., 1997; Shi and Theg, 2010; Su and Li, 2010; Rosano et al., 2011; Inoue et al., 2013; Huang et al., 2016). These chaperones could, in principle, stabilize thylakoid TM proteins in the stroma.

Of these, the chaperonin (Cpn60) is a seemingly unlikely candidate to assist cpSec1-mediated targeting of unfolded proteins. Chaperonins bind non-native proteins and ATP within one heptameric ring of a barrel-shaped tetradecameric oligomer (Hayer-Hartl et al., 2016; Zhao and Liu, 2018). Cpn60 is a heterooligomer of distinct α and β 60-kDa subunits (Musgrove et al., 1987; Nishio et al., 1999). In angiosperms, Cpn60 α further diversified into α1 and α2 isoforms, while Cpn60 β diversified into β1 and β4 isoforms (Zhao and Liu, 2018). Why Cpn60 has so many isoforms remains unclear, but mounting evidence suggests different isoforms specialize in the recognition of distinct substrates (Peng et al., 2011; Ke et al., 2017; Zhao and Liu, 2018). Chaperonins encapsulate bound proteins within one ring in cooperation with a co-chaperonin lid (Hayer-Hartl et al., 2016). ATP hydrolysis within the occupied ring and non-native protein binding to the opposite ring release the encapsulated protein. A released protein may rebind until it achieves a native conformation (Liu et al., 2010; Gruber and Horovitz, 2016; Hayer-Hartl et al., 2016). Chaperonins fold essential proteins in chloroplasts, mitochondria, and bacteria, as does the related group II chaperonin (CCT/TRiC) found in the eukaryotic cytosol (Kerner et al., 2005; Yam et al., 2008; Zhao and Liu, 2018; Bie et al., 2020). Though some biochemical and genetic evidence suggests *E. coli* GroEL (Castanié-Cornet et al., 2014) and mammalian cytosolic CCT/TRiC (Génier et al., 2016) play a role in targeting, all known functions of Cpn60 involve folding stromal proteins.

The nuclear-encoded TM protein Plsp1 localizes to the envelope in meristematic plastids, then increasingly to thylakoids as the chloroplast develops (Shipman and Inoue, 2009). Plsp1 derives from the endosymbiont (Hsu et al., 2011). While Plsp1’s *E. coli* homolog (de Gier et al., 1996; Valent et al., 1998) and presumably its cyanobacterial progenitor insert co-translationally, Plsp1 integrates post-translationally into the envelope by an unknown mechanism and into the thylakoid via the cpSec1 translocon (Endow et al., 2015). During its targeting, Plsp1 transiently resides in the stroma in a 700-kDa complex hypothesized to assist its targeting to the thylakoid (Endow et al., 2015). The size and ATP-mediated dissociation of this complex suggested it represents an interaction between Plsp1 and Cpn60 (Endow et al., 2015). In this work, we demonstrate that Plsp1 interacts with Cpn60 via sequences immediately adjacent to its thylakoid targeting signal and characterize the mechanism by which Cpn60 captures and releases Plsp1 during thylakoid targeting.

## Results

### Plsp1 interacts with Cpn60

Cpn60 migrates in an approximately 700-kDa complex that dissociates upon ATP treatment, the same size and behavior observed of Plsp1’s stromal complex (Hemmingsen and Ellis, 1986; Endow et al., 2015). To confirm that the co-migration of Cpn60 and Plsp1’s stromal complex indeed represented direct interaction between Cpn60 and Plsp1, we determined which stromal proteins Plsp1 binds by an in vitro pulldown assay. Two sequential elution steps tested, first, what was released from binding Plsp1 by ATP treatment, which disrupted Plsp1’s stromal complex (Endow et al., 2015), and, second, what proteins eluted with Plsp1. The Plsp1 variant (mPlsp1_ΔTMD_) used as bait lacks its transit peptide and TMD, which were previously shown to be dispensable for formation of the 700-kDa complex (Endow et al., 2015). As negative controls, we chose the model cpSec1 substrate OE33, which does not interact with Cpn60 (Molik et al., 2001), as well as a bait-less mock pulldown. When incubated with freshly prepared SE, Plsp1 pulled down Cpn60. This Cpn60 was partially released by ATP treatment when Plsp1, but not OE33, was the pulldown bait (Fig. 1A, lane 6). More Cpn60 eluted with Plsp1 (Fig. 1A, lane 8). In contrast, OE33 eluted with considerably less than 1% of input Cpn60 (Fig. 1A, compare lanes 1 and 8). These results indicate that Cpn60 interacted directly and specifically with Plsp1.

**Figure 1.**
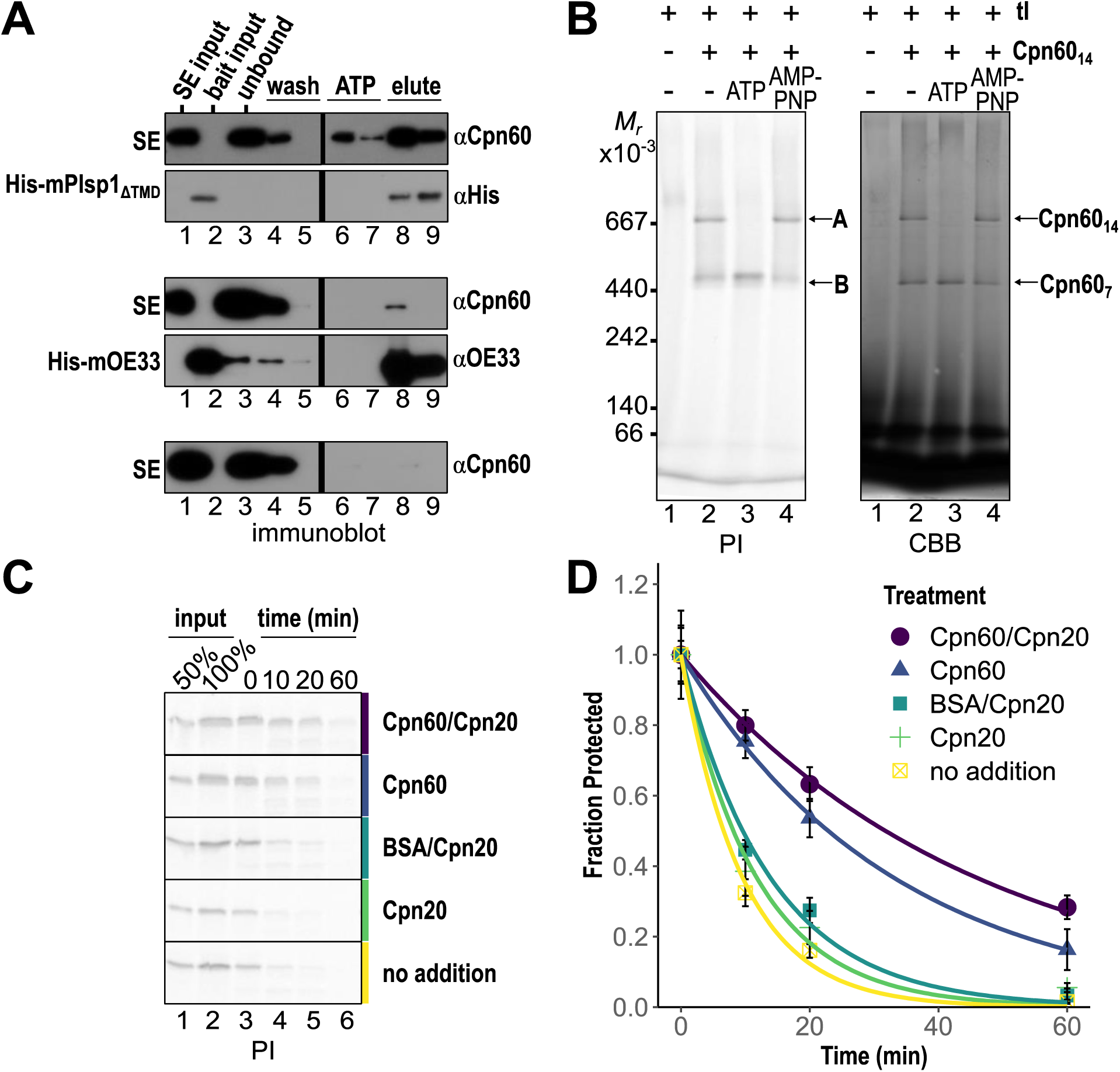
Plsp1 directly interacts with Cpn60 α1 and β1. (A) In vitro pulldown of Cpn60 from stromal extract (SE) with His-tagged bait. 20 μg bait proteins were incubated with SE equivalent to 250 μg chlorophyll (Chl), then bound to affinity resin, washed, and sequentially treated with 10 mM ATP and 300 mM imidazole. Representative immunoblots of three independent experiments show 1% of SE input, 2.5% of bait input, 1% of unbound, and 25% of two washes, ATP treatment (ATP), and imidazole elution (elute%). Black lines mark removed lanes (three additional washes). Cpn60 was detected with αHsp60; His-mPlsp1_ΔTMD_ with αHis, and His-mOE33 with αOE33. (B) Cpn60 tetradecamer binding assay. In vitro-translated, radiolabeled mPlsp1 was incubated with 250 nM Cpn60 tetradecamer (Cpn60_14_) for 30 min at room temperature. 10 mM ATP or AMP-PNP were added (where indicated) for an additional 10 min incubation. The binding assay was separated on 4-14% BN-PAGE. Radiolabeled proteins were visualized by autoradiography and phosphorimaging (left, PI), and Cpn60 oligomers were visualized by Coomassie staining (right, CBB). Cpn60_14_ and Cpn60_7_ indicate the sizes of tetradecameric and heptameric Cpn60 oligomers, respectively. (C) Proteinase K protection assay of Cpn60-mediated substrate encapsulation. In vitro-translated, radiolabeled Plsp1 was incubated with Cpn60_14_ (0.8 μM), BSA (11.2 μM, equivalent to Cpn60 protomer concentration), or buffer for 30 min at room temperature to allow Plsp1 to bind. ADP and, where indicated, Cpn20_4_ (2 μM), were added, followed by 10 mins of further incubation. Assay was initiated with the addition of 1.2 μg/ml Proteinase K, and aliquots were removed at 0, 10, 20, and 60 minutes and quenched with PMSF. After separation on SDS-PAGE, radiolabeled Plsp1 was visualized by autoradiography. Gel phosphorimages (PI) shown are representative of N=6 (no addition) or N=7 (all other treatments) independent digestions. (D) Quantification of (C). Protection at each time point for each treatment was quantified relative to the 0-minute time point. Points represent mean; error bars represent ± 1 SD. Lines represent single-exponential curves fit to data.

The specific Cpn60 isoforms and other stromal proteins pulled down by Plsp1 were identified by LC-MS/MS. Proteomic analysis demonstrated that the proteins eluted with Plsp1 (Fig. 1A, lane 8) included Cpn60α1 and three β1 isoforms (Table 1). The minor Cpn60α2 and β4 isoforms and the co-chaperonins Cpn20 and Cpn10 were not pulled down to an appreciable degree (Table 1). An additional stromal chaperone, Hsp70, was also found in the Plsp1 pulldown. The Hsp70 log_2_ fold change of 4.7 represented a statistically significant 25-fold enrichment in the pulldown compared to the control (Table 1). However, Cpn60 was pulled down in near stoichiometric amounts, but Hsp70 was only 1.6% of the Cpn60 pulled down (Table 1, Supplemental Dataset 1). The bulk of stromal Hsp70 did not co-migrate with Plsp1’s 700-kDa complex (Endow et al., 2015). These data suggest that while Hsp70 can interact with Plsp1, Hsp70 is unlikely to be responsible for the observed stromal complex. Nine additional proteins were pulled down by Plsp1; however, these proteins were minimally abundant as compared to Cpn60 (Supplemental Data Set 1). From this in vitro pulldown, we conclude that Plsp1 directly interacts with Cpn60α1 and β1.

**Table 1.**
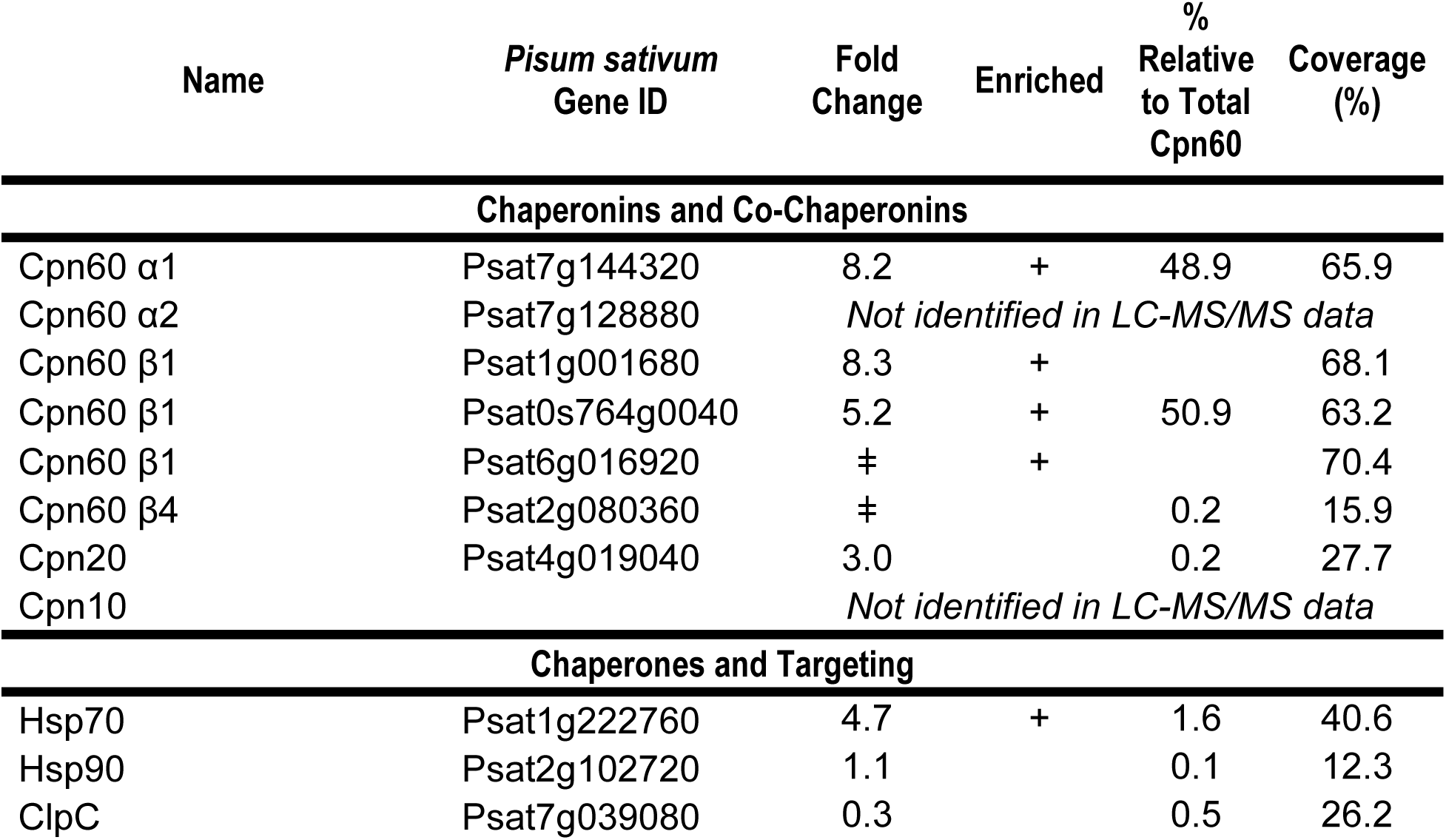
Stromal chaperones pulled down by Plsp1. LC-MS/MS analysis of proteins pulled down by Plsp1 (Fig. 1A, lane 8). The average fold change between pulldown (mPlsp1_ΔTMD_ bait) and control (SE-only mock) was calculated from the log_2_ transformed ratio of intensity-based absolute quantification (iBAQ) values for each protein; ‡ indicates detection solely in the pulldown. Enriched (+) indicates proteins which were significantly more abundant in the pulldown than the control; enrichment was determined by Student’s t-test (p < 0.05) with correction for multiple comparisons. The % relative to total Cpn60 was calculated using the total iBAQ value for all detected Cpn60 isoforms. The three highly similar Cpn60 β1 isoforms were combined for the % relative to total Cpn60 calculation due to difficulty distinguishing isoforms. Nine additional enriched proteins, none of which amounted to greater than 0.5% total Cpn60, were excluded from this table due to their low abundance; full data for all detected proteins are in Supplementary Data Set 1. N = 2 independent experiments.

We tested if isolated Cpn60 tetradecamers (Cpn60_14_) and in vitro translated, radiolabeled Plsp1 were sufficient to reconstitute the 700-kDa complex. Cpn60_14_ prepared from *Pisum sativum* Cpn60α1 and β1 (Dickson et al., 2000) was incubated with radiolabeled Plsp1 and examined by BN-PAGE. Two prominent bands of radiolabeled Plsp1 were detected, migrating just above the 669- and 440-kDa markers (Fig. 1B, labeled A and B). Band A is the same size as Plsp1’s stromal 700-kDa complex and co-migrates with the Cpn60_14_ band detected by Coomassie staining. Band B corresponds approximately in size to a single ring Cpn60 oligomer (Cpn60_7_) and co-migrates with a fainter Coomassie-staining band. Mitochondrial Hsp60 has active single- and double-ring forms (Weiss et al., 2016), but an active Cpn60_7_ has not been previously described. Band A disappeared in the presence of hydrolysable ATP, consistent with the observed behavior of the stromal 700-kDa complex (Endow et al., 2015). 10 mM ATP disrupted Band A (Fig. 1B, lane 3), while 10 mM AMP-PNP had no effect (lane 4). Band B increased in intensity upon 10 mM ATP treatment, as did its co-migrating Cpn60_7_ band. We conclude that Cpn60α1, Cpn60β1, and Plsp1 interact to form a complex equal in size and similar in behavior to that observed on import into chloroplasts.

Chaperonin interaction conveys protection of its substrates to Proteinase K, especially in the presence of co-chaperonin, as the substrate is encapsulated within the chaperonin’s cavity beneath the co-chaperonin lid (Weissman et al., 1995; Mao et al., 2015). We used this property to further validate that Plsp1 interacts with Cpn60. Radiolabeled Plsp1 was incubated with Cpn60_14_, BSA, or buffer to allow Plsp1 to bind, then supplemented with Cpn20 (where indicated). BSA incubation served as a control for slow degradation under high protein load. After the addition of Proteinase K, samples were removed at 0, 10, 20, and 60 min, and the fraction of Plsp1 remaining assessed (Fig. 1C-D). At all time points, Plsp1 exhibited partial resistance to degradation in the presence of Cpn60 as compared to all controls (Fig. 1C). Cpn20 alone did not protect Plsp1 but had a small additive effect on the protection due to Cpn60 under these conditions (Fig. 1D). The higher level of protection of Plsp1 in the presence of Cpn60_14_ than in the presence of equimolar BSA demonstrated that the protection of Plsp1 was due to Cpn60 interaction specifically, not the protein concentration of the reaction (Fig. 1C-D). The presence or absence of Plsp1’s transit peptide did not alter its protection by Cpn60 (Supplemental Fig. 1). Collectively, these data support the conclusion that Cpn60 encapsulates Plsp1.

### The dissociation of the 700-kDa complex correlates with increased membrane integration

Plsp1 transiently resides in the stroma prior to its integration into thylakoids by cpSec1, and interaction with Cpn60 may be an intermediate step in Plsp1’s targeting (Endow et al., 2015). We expanded the import-chase assay from our previous study to examine how Plsp1’s interaction with Cpn60 correlates with its membrane integration. To generate a pulse of Plsp1, Plsp1 was imported for 10 minutes, then unincorporated Plsp1 was degraded with membrane-impermeable thermolysin. Chloroplasts were returned to import conditions for an additional 0, 10, or 30 minutes of chase incubation. The soluble fractions at each chase time point, visualized by BN-PAGE, contained three distinguishable radiolabeled bands, approximately 180-kDa, 550-kDa, and 700-kDa in size (Fig. 2A). The 700-kDa region contained all major and minor isoforms of Cpn60, and Cpn60 was its most abundant complex (Fig. 2B). The relative abundance of Cpn60 isoforms suggest an α1_6_β1_8_ stoichiometry, as reported for *Chlamydomonas* (Bai et al., 2015; Zhao et al., 2019), with α2 and β4 representing about 0.2 and 0.4 % of the total Cpn60, respectively (Supplemental Data Set 2). The intensity of the 700-kDa complex decreased over chase-time (Fig. 2A), consistent with the observed overall decline in Plsp1 detected in the soluble fraction (Fig. 2C). The amount of Plsp1 in the membrane fraction increased, and treatment of the membranes with thermolysin showed that Plsp1’s degradation product (dp1), which indicates topologically-correct integration, also increased over chase-time (Fig. 2C), consistent with previous results (Endow et al., 2015). The total level of imported Plsp1 stayed consistent throughout the chase, indicating that Plsp1 is not degraded (Fig. 2C). The decline in the 700-kDa complex therefore correlated with the increased membrane integration (Fig. 2D). The behavior of the 700-kDa complex is consistent with it serving as an early intermediate in the targeting of Plsp1, as Plsp1 dissociates from Cpn60 prior to its maximal membrane integration.

**Figure 2.**
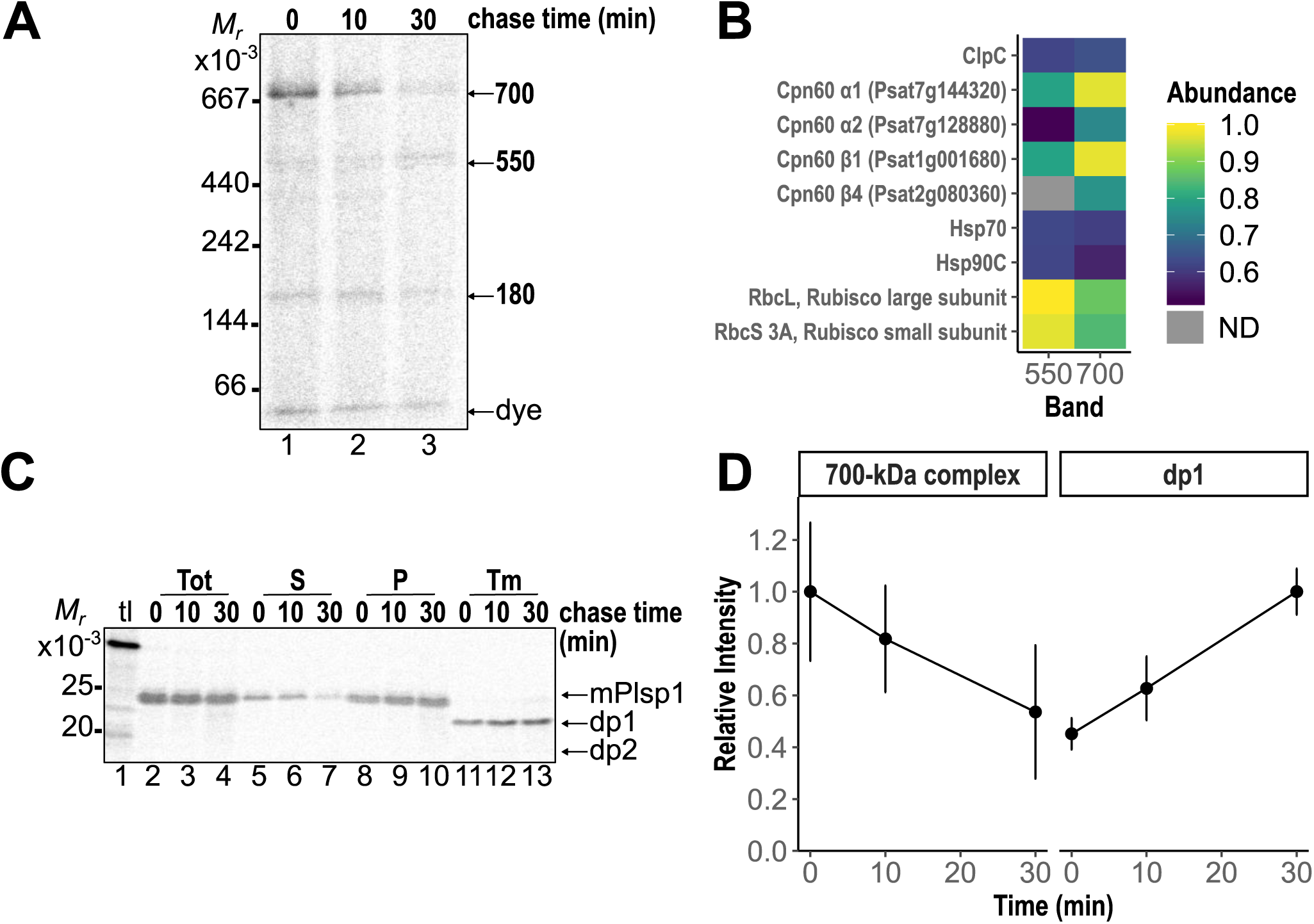
The level of Plsp1 interacting with Cpn60 decreases over chase-time. (A) Import-chase assay followed by BN-PAGE analysis. In vitro-translated, radiolabeled Plsp1 was imported into illuminated (50 μE/m^2^s) chloroplasts with 3 mM ATP for 10 min, followed by treatment with thermolysin for 30 min on ice in the dark to remove unincorporated precursor. The protease was quenched, and intact chloroplasts were reisolated and incubated in 50 μE/m^2^s light with 3 mM ATP for 0, 10, or 30 min. Shown is a representative phosphorimage of the equivalent of 6 μg chlorophyll (Chl) of soluble fraction from each chase time-point analyzed by BN-PAGE and autoradiography. Import and chase reactions contained 1 mM lincomycin to suppress chloroplast translation of RbcL. (B) Subset of proteins identified in LC-MS/MS analysis of the 550-kDa and 700-kDa regions. The 700-kDa band was excised from the soluble fraction after 10 min import; the 550-kDa band was excised from the soluble fraction after 10 min import and 30 min chase. iBAQ (intensity-based absolute quantification) values from each excised band were normalized to the median. To facilitate easier comparison, abundance values for each protein were divided by the average abundance of RbcL in the 550-kDa band (the most abundant protein in any band). Colors indicate relative abundance; gray indicates not detected (ND). Data represent two independent experiments for each band, and independent experiments were performed in triplicate (700-kDa) or duplicate (550-kDa). For all identified proteins, see Supplemental Data Set 2. (C) Import-chase assay as described in (A), followed by SDS-PAGE analysis of chloroplast subfractions. Representative phosphorimage of the equivalent of 3 μg Chl of total import (Tot), soluble (S), membrane-associated (P), and thermolysin-resistant (Tm) fractions of each chase time-point. Equal Chl equivalents were run on SDS-PAGE alongside 10% input translation (tl). Sizes of mature (m), dp1, and dp2 Plsp1 are indicated. (D) The intensity of the 700-kDa complex at each chase time point was quantified relative to its peak (t=0 min). The amount of thermolysin-protected dp1 at each chase time point was quantified relative to its peak (t=30 min). Import-chase assays included in this quantification were performed without lincomycin and analyzed stromal proteins equivalent to 3 μg Chl on both BN-PAGE and SDS-PAGE. Points represent mean (N=10 for 700-kDa complex; N=7 for dp1); error bars are ± 1 SD

The 180-kDa and 550-kDa bands were less prominent than the 700-kDa Cpn60 complex (Fig. 2A). The 180-kDa band did not increase or decrease consistently through the chase (Supplemental Fig. 2) and, as such, was not further characterized. The 550-kDa band increased over chase-time (Fig. 2A, Supplemental Fig. 2). The 550-kDa region contained primarily RbcL and RbcS as well as a moderate amount of the major Cpn60 isoforms (Fig. 2B). The Rubisco holoenzyme and the single ring Cpn60 complex observed with isolated oligomers above (Fig. 1B) both migrate at this size. The increase in the 550-kDa band was largely suppressed when chloroplast translation was inhibited with lincomycin (Supplemental Fig. 3A-B). Though Plsp1 may contribute to the 550-kDa signal, background synthesis of radiolabeled RbcL appears responsible for its increase. These results indicate that the 180-kDa and 550-kDa bands are not intermediates in Plsp1 targeting.

**Figure 3.**
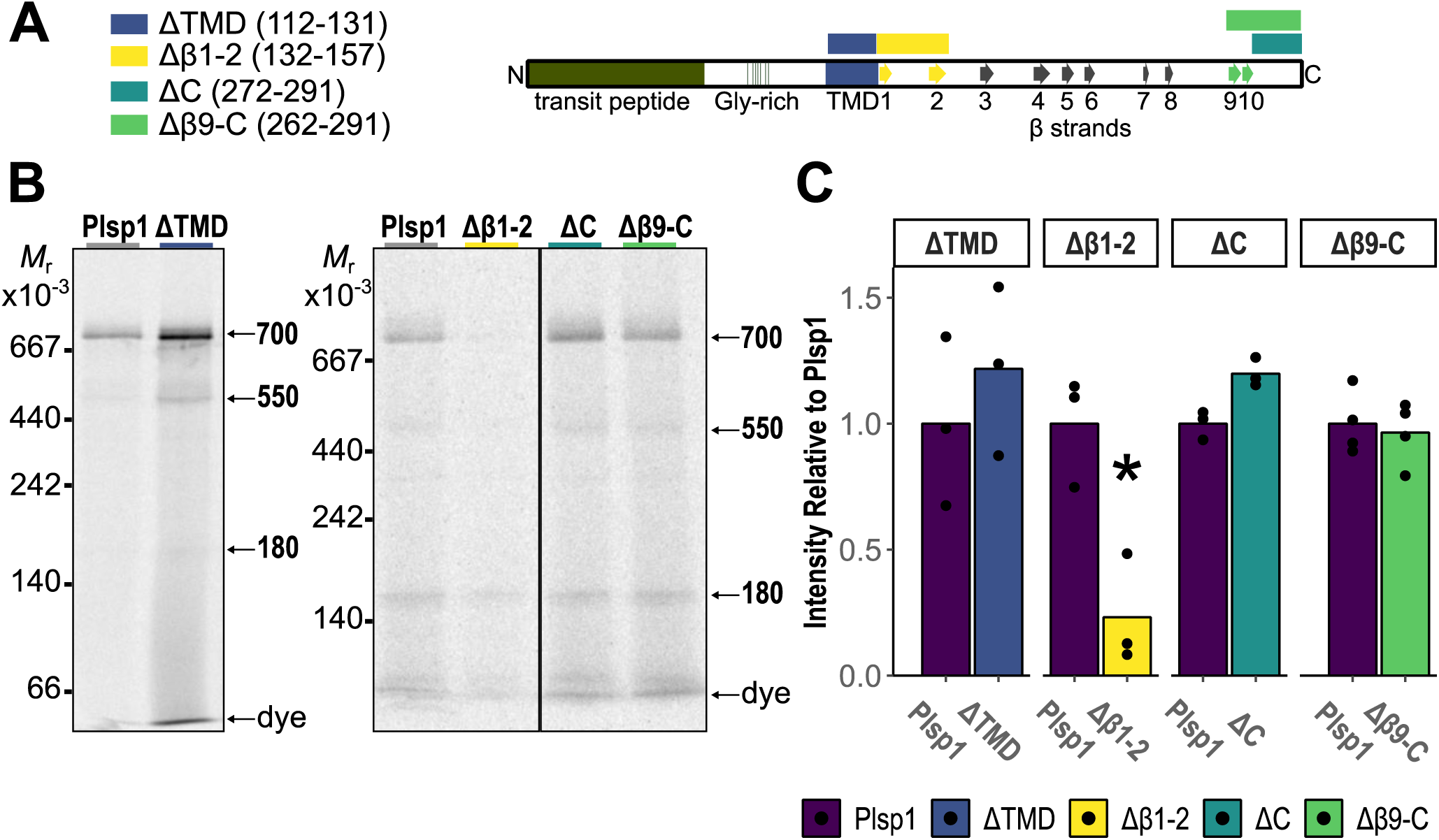
β1-2 is necessary for efficient interaction with Cpn60. (A) Sequence diagram of full-length Plsp1 with predicted transmembrane domain (TMD) and β sheets indicated. β1-2 and β9-10 correspond by homology to the hydrophobic membrane association surface in the *E. coli* leader peptidase crystal structure. Colored bars indicate residues removed for deletion constructs: ΔTMD (112-131, blue), Δβ1-2 (132-157, yellow), ΔC (272-291, teal), and Δβ9-C (262-291, green). (B) Import assay followed by BN-PAGE analysis. After 10 min import of full-length Plsp1 or deletion constructs, intact chloroplasts were lysed hypotonically. The soluble fraction was collected by centrifugation, separated on 4-14% BN-PAGE, and visualized by autoradiography. A sample of intact chloroplasts was analyzed directly by SDS-PAGE to confirm import and calculate import efficiency. Black lines on gel indicate lane removal. N = 3 for each deletion construct. (C) To quantify the 700-kDa complex, constructs were analyzed pairwise with their full length Plsp1 control. Densitometry signals for each deletion construct and the full length Plsp1 control were normalized to account for different numbers of methionine residues and import efficiency. Signals from an individual experiment were normalized to the mean of both signals from that experiment; normalized signals for full length Plsp1 and a deletion construct were then divided by the mean of signals for full length Plsp1 for all replicates. Significant differences (p<0.05, *) between mean full length and deletion construct were calculated by t-test.

### Cpn60 and cpSec1 bind adjacent Plsp1 sequences

The correlation between increasing membrane association and decreasing Cpn60 interaction suggests that Cpn60 hands Plsp1 off to cpSec1. Our ability to investigate this hypothesis with genetic approaches was limited because Cpn60 is essential (Zhao and Liu, 2018). We therefore interrogated the Plsp1 domains necessary for interaction with Cpn60 and cpSec1 and the effect of specific stromal factors on integration with in vitro targeting assays.

If Cpn60 binds a specific Plsp1 sequence, we predicted that removing the sequence would impair formation of the 700-kDa complex during import. We cloned Plsp1 variants that deleted the transmembrane domain (TMD), the β strands (β 1-2 and 9-10) predicted to form the membrane association surface, and the extreme C-terminus (Fig. 3A) (Endow et al., 2015; McKinnon et al., 2020). These regions all contain hydrophobic segments likely to bind chaperones. After each deletion construct was imported, we isolated the stroma for BN-PAGE analysis and compared the intensity of the 700-kDa complex to a full-length Plsp1 control. We found that Δβ1-2 exhibited a consistent 80% reduction in the intensity of the 700-kDa band, indicating β1-2 is necessary for interaction with Cpn60 (Fig. 3B-C). In contrast, loss of Plsp1’s TMD or C-terminus did not alter the 700-kDa complex (Fig. 3B-C). The 550-kDa and 180-kDa bands were not reduced for any deletion construct (Supplemental Fig. 4A-B). These results indicate that the β strands immediately adjacent to the TMD are important for the interaction of Plsp1 with Cpn60.

**Figure 4.**
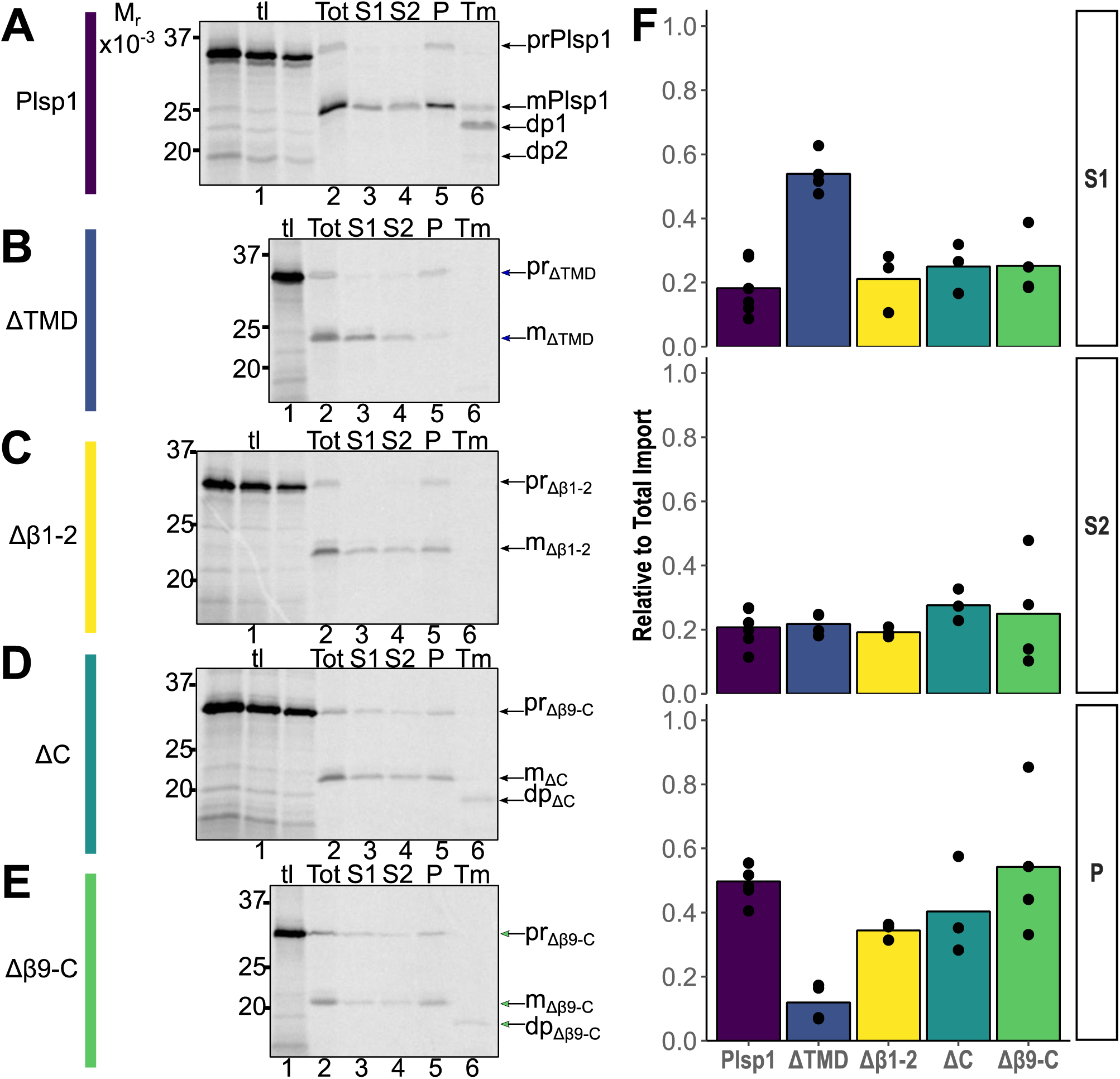
TMD and β1-2 are necessary for membrane integration. (A) Subfractionation and integration analysis of full-length Plsp1. After 10-min import, intact chloroplasts were separated into three samples to analyze total import (Tot), subfractionation, and integration (Tm). The first sample (Tot) was pelleted and analyzed directly. The second sample was lysed hypotonically, and the soluble material (S1) was collected by centrifugation. Membranes were subsequently washed with 0.1M Na_2_CO_3_ to extract peripheral membrane proteins. The alkaline-extracted fraction (S2) was separated from the alkaline-resistant membranes (P) by centrifugation. The third sample (Tm) was lysed hypotonically and treated with thermolysin. Equal chlorophyll (Chl) equivalents from each sample were separated on SDS-PAGE alongside 10% translation product (tl). The precursor (pr), mature (m), and thermolysin degradation products of successfully targeted (dp1) and mistargeted (dp2) Plsp1 are indicated. (B) ΔTMD, analyzed as in (A). (C) Δβ1-2, analyzed as in (A). (D) ΔC, analyzed as in (A). (E) Δβ9-C, analyzed as in (A). (F) The amount of mature protein relative to the total import (Tot) was calculated for S1, S2, and P. Bars represent mean; points represent measurements from at least three independent experiments.

Plsp1 lacks a cleavable thylakoid targeting signal, so cpSec1 must recognize a domain of Plsp1’s mature sequence. Deleting that domain may cause defects in targeting, which we visualized by comparing the subfractionation patterns of full-length Plsp1 and the deletion constructs after a 10-min import (Fig. 4A-E). We quantified how much mature protein, relative to the total imported mature protein, fractionated with soluble proteins (S1), alkaline-extracted peripheral proteins (S2), and alkaline-resistant membrane proteins (P) (Fig. 4F). Full-length Plsp1 distributed to S1, S2, and P in a 1:1:2 ratio, with approximately 18% in S1 (Fig. 4A). The ΔTMD construct stalled in the stroma, with 54% found in S1 – a three-fold increase over full-length Plsp1 – and only traces associating with the alkaline-resistant pellet (Fig. 4B). We previously showed Plsp1’s TMD was necessary for cpSec1-dependent integration into isolated thylakoid membranes (Endow et al., 2015). Together, these data indicate that Plsp1’s TMD comprises its thylakoid targeting signal. Despite altered association with the 700-kDa complex, Δβ1-2 distributed among subfractions indistinguishably from full-length Plsp1 (Fig. 4C), as did ΔC and Δβ9-C (Fig. 4D-E). These fractionation patterns indicate all deletions except ΔTMD transit the stroma and reach the thylakoid membrane.

Full integration of Plsp1 into thylakoids results in the characteristic degradation product (dp1) after treatment with the membrane-impermeable protease thermolysin (Fig. 4A, lane 6). Neither ΔTMD nor Δβ1-2 yielded such a degradation product after import for 10 min (Fig. 4B-C, lane 6). Extended 30-min import of Δβ1-2 yielded a faint degradation product that was only about 10% of dp1 from full-length Plsp1 (Supplemental Fig. 5). ΔC and Δβ9-C both produced a dp1-equivalent degradation product (Fig. 4D-E, lane 6), suggesting that the C-terminus of Plsp1 is not necessary for successful insertion. ΔTMD did not associate with the thylakoid membrane (Fig. 4B), hence its impeded integration. The inability of Δβ1-2 to bind Cpn60 may result in Δβ1-2 assuming a conformation incompetent for insertion by cpSec1. The β1-2 sequence is also adjacent to the TMD that acts as Plsp1’s thylakoid targeting signal (Fig. 3A), and its deletion may remove part of the sequence that cpSec1 requires to insert Plsp1. The adjacent TMD and β1-2 regions control membrane integration. Cpn60 and cpSec1 interact with adjoining sequences of Plsp1.

**Figure 5.**
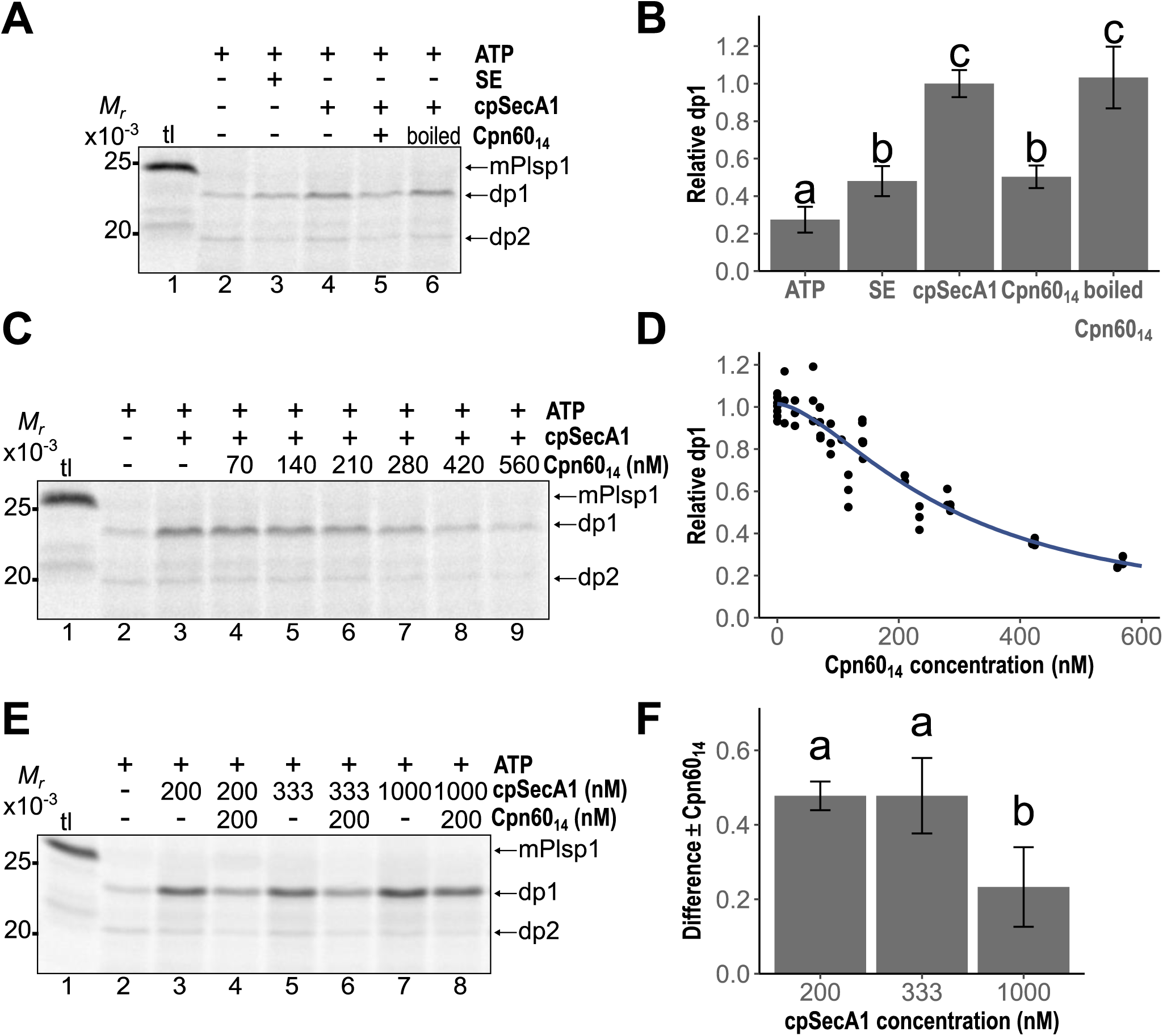
The balance of Cpn60_14_ to cpSecA1 influences integration. (A) In vitro transport assay. Radiolabeled Plsp1 was incubated with thylakoids in the presence of 5 mM ATP (lanes 2-6), stromal extract (SE, lane 3), 200 nM cpSecA1 (lanes 4-6), and 200 nM Cpn60_14_ (lane 5; lane 6, boiled) for 30 min in 90 μE/m^2^s light. Thylakoids were treated with thermolysin, and equal chlorophyll (Chl) equivalents were separated by SDS-PAGE alongside 5% translation product (tl) and visualized by autoradiography. The sizes of mature (mPlsp1) and thermolysin degradation products (dp1, dp2) are indicated. Shown is a representative phosphorimage of three independent experiments performed in triplicate. (B) Quantification of dp1 from (A) relative to dp1 in the presence of ATP and cpSecA1 (A, lane 4). Bars represent mean; error bars represent ± 1 SD. Means of the treatments with the same letter do not differ significantly based on Tukey HSD for α = 0.05 (C) In vitro transport assay with different Cpn60_14_ concentrations. Radiolabeled Plsp1 was incubated with thylakoids in the presence of 5 mM ATP (lanes 2-9), 200 nM cpSecA1 (lanes 3-9), and the indicated concentrations of Cpn60_14_ (lanes 4-9) for 30 min in 90 μE/m^2^s light. Thylakoids were treated with thermolysin and analyzed as in (A). Shown is a representative phosphorimage of three independent experiments, each in triplicate; exact concentrations of Cpn60_14_ varied between independent experiments (D) Quantification of dp1 from (C) relative to dp1 in the presence of ATP and cpSecA1 (C, lane 3). All replicates are plotted as black points, with a dose-dependence curve (Gadagkar and Call, 2015) fit to illustrate the relationship between Cpn60_14_ concentration and integration. (E) In vitro transport assay with different cpSecA1 concentrations. Radiolabeled Plsp1 was incubated with thylakoids in the presence of 5 mM ATP (lanes 2-8) and 200 nM (lanes 3-4), 333 nM (lanes 5-6), or 1000 nM (lanes 7-8) cpSecA1 for 30 min in 90 μE/m^2^s light. In lanes 4, 6, and 8, 200 nM Cpn60_14_ was included. Thylakoids were treated with thermolysin and analyzed as in (A). Shown is a representative phosphorimage of three independent experiments performed in triplicate. (F) Quantification of the difference in dp1 (E) in the presence and absence of 200 nM Cpn60_14_. dp1 was quantified relative to dp1 in the presence of ATP and 200 nM cpSecA1 (E, lane 3), and then the difference between + Cpn60_14_ and – Cpn60_14_ integration was calculated for each concentration of cpSecA1. Bars represent mean difference; error bars represent ± 1 SD. Means of the differences with the same letter do not differ significantly based on Tukey HSD for α = 0.05.

### The balance of Cpn60 and cpSec1 controls Plsp1 integration

With increased mechanistic insight into how Cpn60 and cpSec1 recognize Plsp1, we next asked how Cpn60 and cpSecA1, the extrinsic cpSec1 motor, influenced integration of Plsp1 into isolated thylakoids. Cpn60 and cpSecA1 do not interact in the stroma (Supplemental Fig. 6). We previously demonstrated that Plsp1 is integrated by stromal extract (SE)-independent (Fig. 5A, lane 2) and SE-dependent (Fig 5A, lane 3) mechanisms using an in vitro transport assay (Endow et al., 2015). Both mechanisms inserted Plsp1 correctly, resulting in dp1 after thermolysin treatment, and both mechanisms produced in a second degradation product (dp2). Data suggest dp2 represents mistargeted association with the stromal face of the thylakoid (Endow et al., 2015). SE-independent insertion displayed a higher proportion of dp2 (Fig. 5A, lanes 2), indicating more mistargeting. The SE-dependent insertion is mediated by cpSec1 (Endow et al., 2015), and, as previously shown, adding cpSecA1 to the transport assay stimulated dp1 (Fig. 5A, lane 4). To test the effect of Cpn60 on integration, we performed the in vitro transport assay in the presence of equimolar Cpn60_14_ and cpSecA1. These conditions decreased dp1 formation by 50% (Fig. 5A, lane 5) compared to the addition of cpSecA1 alone (lane 4). When the added Cpn60 was inactivated by boiling, dp1 did not decrease (Fig. 5A, lane 6). Thus, active Cpn60 reduced integration driven by equimolar cpSecA1.

**Figure 6.**
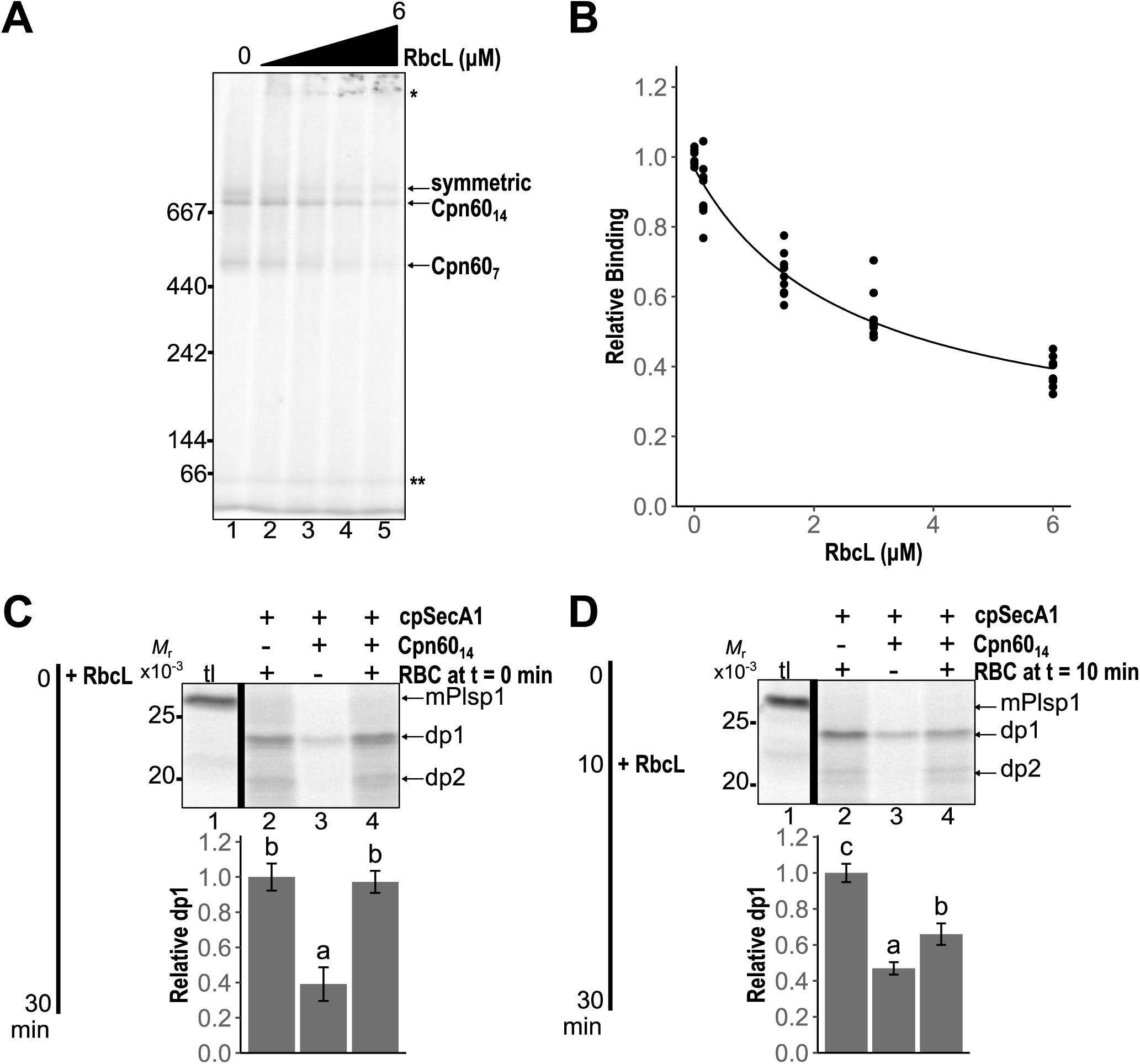
Cpn60-bound Plsp1 is released by competition with RbcL. (A) Competition binding assay. In vitro-translated, radiolabeled mPlsp1 was incubated with 200 nM Cpn60_14_ for 5 min at room temperature, then chilled on ice for 5 min before guanidine-denatured RbcL was diluted 36-fold into the reaction. Final concentrations were 0, 0.15, 1.5, 3, or 6 μM RbcL (lanes 1-5, respectively). Reactions were transferred to room temperature, then analyzed by BN-PAGE and autoradiography. Shown is a representative phosphorimage of three independent experiments performed in triplicate. (B) The intensity of the Cpn60_14_ band indicated in (A) was quantified relative to addition of 0 μM RbcL. Points represent values from independent experiments; line represents a dose-dependent response curve (Gadagkar and Call, 2015) fit to points. (C) In vitro transport with RbcL added at t = 0 min. Radiolabeled Plsp1 was mixed with thylakoids in the presence of 5 mM ATP (lanes 2-5), 200 nM cpSecA1 (lanes 2-5), and, where indicated, 200 nM Cpn60_14_ (lanes 4-5) on ice. Guanidine-denatured RbcL or the equivalent buffer control were diluted 36-fold into transport reaction (3 μM, final), which was illuminated (90 μE/m^2^s) at room temperature for 30 min. Thylakoids were treated with thermolysin and analyzed as in Fig. 5A. 5% translation input (tl) and mature, dp1, and dp2 sizes are indicated. dp1 was quantified for each treatment and normalized to transport with cpSecA1 and RbcL (lane 3). Bars represent mean; error bars represent ± 1 SD. Means of the treatments with the same letter do not differ significantly based on Tukey HSD for α = 0.05. N = 3 independent experiments, performed in triplicate (D) In vitro transport with RbcL added at t = 10 min. Radiolabeled Plsp1 was incubated with thylakoids in the presence of 5 mM ATP, 200 nM cpSecA1, and, where indicated, 200 nM Cpn60_14_ (lanes 3, 5) for 10 min in 90 μE/m^2^s light at room temperature. Guanidine-denatured RbcL or the equivalent buffer control were diluted 36-fold into transport (3 μM, final) and incubated for 20 min further in 90 μE/m^2^s light at room temperature. Thylakoids were treated with thermolysin and analyzed as in Fig. 5A. dp1 was quantified and normalized as in (D). Bars represent mean; error bars represent ±1 SD. Means of the treatments with the same letter do not differ significantly based on Tukey HSD for α = 0.05. N = 3 independent experiments, performed in triplicate.

Cpn60_14_ and cpSecA1 are not equimolar in the stroma. We estimated the concentration of Cpn60 α1 and cpSecA1 in SE by immunoblotting with recombinant protein standards (Supplemental Fig. 7). With 95% confidence, there were 26 to 37 ng Cpn60 α1 and 6 to 8 ng cpSecA1 per μg stromal protein. The calculated Cpn60_14_ concentration was 24 μM, in rough agreement with the reported concentration of 13 μM (Musgrove et al., 1987). The calculated cpSecA1 concentration was 15 μM. In *E. coli*, it is disputed whether SecA functions as a monomer or dimer (Cranford-Smith and Huber, 2018); in chloroplasts, its migration suggests a dimer (Nakai et al., 1994; Takabayashi et al., 2017). The approximate 3 Cpn60_14_ : 2 cpSecA1 molar ratio will further vary based on Cpn60 occupancy and transient local concentration differences. We therefore tested the effect of a range of Cpn60_14_ and cpSecA1 concentrations in the Plsp1 transport assay. At 200 nM cpSecA1, the reduction in dp1 increased as higher concentrations of Cpn60_14_ were added (Fig. 5C). Up to 140 nM Cpn60_14_ had little effect on dp1, but 560 nM Cpn60_14_, the maximum added, reduced dp1 by 77% (Fig. 5D). The concentration of Cpn60_14_ which reduced dp1 by 50% was calculated to be 290 nM. The reduction was also specific to Cpn60 as equimolar BSA did not affect dp1 (Supplemental Fig. 8). While 200 nM Cpn60_14_ reduced dp1 by 48% at equimolar cpSecA1, the same concentration of Cpn60_14_ only reduced dp1 by 23% at 1 μM cpSecA1 (Fig. 5E-F). Increasing cpSecA1 stimulates integration and compensates for the Cpn60-mediated reduction. The balance of Cpn60 to cpSecA1 controls the extent to which Plsp1 is integrated.

**Figure 7.**
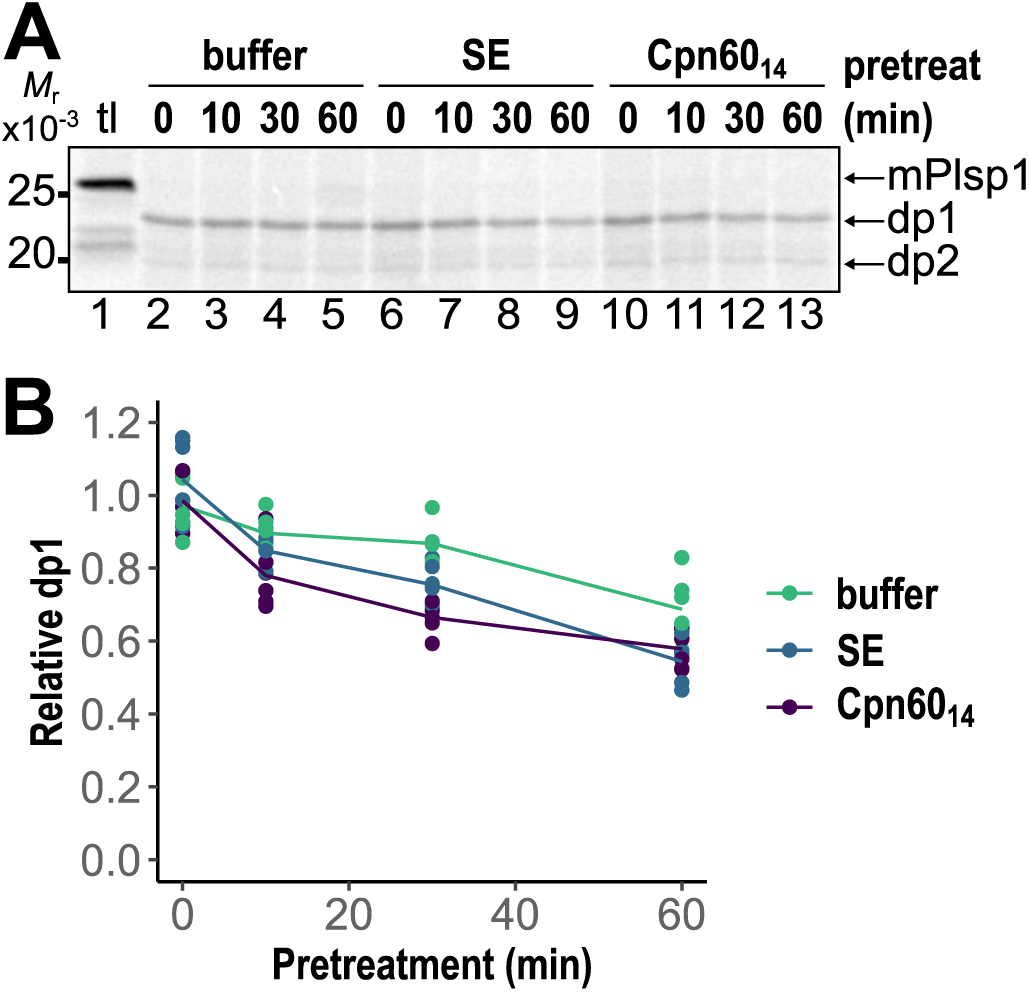
Neither Cpn60 nor other stromal chaperones actively maintain Plsp1’s transport competence. (A) In vitro translated, radiolabeled Plsp1 was pretreated with buffer, stromal extract (SE), or 200 nM Cpn60_14_ for 0, 10, 30, or 60 min at room temperature. Pretreatments were adjusted such that all reactions contained final concentrations of 40 nM Cpn60_14_ and 0.2 mg/ml chlorophyll equivalent SE, then incubated with thylakoids in the presence of 5 mM ATP and 200 nM cpSecA1 for 30 min in 90 μE/m^2^s light. Thylakoids were treated with thermolysin and analyzed by SDS-PAGE alongside 5% input translation (tl). Shown is a representative phosphorimage of three independent experiments performed in duplicate. (B) Quantification of dp1 from (A) relative to the mean of all t = 0 min pretreatment samples (lanes 2, 6, 10). Points are all replicates, colored by treatment; lines are drawn to assist visualization.

### Plsp1 displaced from Cpn60 is accessible to cpSec1

To exchange Plsp1 between Cpn60 and cpSec1, Plsp1 must ultimately dissociate from Cpn60. Non-native substrates can rapidly rebind chaperonins after release; however, substrates with higher dependence on chaperonins for folding will out-compete less stringent substrates (Kerner et al., 2005). To determine if Cpn60-bound Plsp1 releases and rebinds, we pre-bound radiolabeled mature Plsp1 to Cpn60, then added an excess of chemically denatured Rubisco as a competitor (Fig. 6A). RbcL is an obligate Cpn60 substrate (Aigner et al., 2017). Higher concentrations of RbcL increasingly displaced the radiolabeled Plsp1 from binding Cpn60 (Fig. 6A). The maximum RbcL added, a 30-fold molar excess over Cpn60_14_, reduced the binding of Plsp1 to 38% of mock RbcL addition (Fig. 6B). Free Plsp1 did not increase at the high concentrations of added RbcL (Fig. 6A, indicated by **), but instead, more radiolabeled protein failed to enter the gel entirely (Fig. 6A, indicated by *). During these experiments, a minor proportion of Cpn60 migrated slightly above the 700-kDa Cpn60_14_ band, which we attribute to substrate occupying both rings in a symmetrical complex (Fig. 6A) (Weiss et al., 2016; Bigman and Horovitz, 2019). From this experiment, we conclude that non-native RbcL can out-compete Plsp1 from rebinding Cpn60.

The release of Plsp1 from Cpn60 should allow its recognition by cpSec1 for membrane integration (Fig. 6A). We tested if the addition of RbcL during the in vitro Plsp1 transport assay alleviated the competition by Cpn60. Plsp1 integration proceeds linearly for approximately 10 minutes, then continues to slowly increase during a 30-min transport. We added 3 μM denatured RbcL at t = 0 min to initiate the assay (Fig. 6D) or at t = 10 min into the Plsp1 transport reaction (Fig. 6E). When RbcL was added to transport reactions at t = 0 min, the amount of dp1 ultimately produced was the same in the presence or absence of Cpn60 (Fig. 6D, compare lanes 3 and 5). The integration of Plsp1 was fully restored by competitive RbcL. When RbcL was added to transport reactions at t = 10 min, the amount of dp1 in the presence of Cpn60 increased but did not fully recover to the level of the control (Fig. 6E). RbcL displacement was less effective at room temperature than on ice (Supplemental Fig. 9, Fig. 6B), which might partially account for the incomplete restoration of integration. From these results, we conclude that Plsp1 released from Cpn60 is accessible to cpSec1.

When cpSRP43 or cytosolic Hsp70 provide chaperone assistance to membrane-targeted substrates, they actively maintain their substrates in an unfolded, insertion-competent conformation (Payan and Cline, 1991; Jaru-Ampornpan et al., 2010; Cho and Shan, 2018). We modified an assay used to demonstrate stromal extract (SE) maintained the transport-competence of LHCP over 1 hour of pretreatment to test the effect of stromal chaperones and of Cpn60 on the ability of Plsp1 to integrate (Payan and Cline, 1991). In our experiments, pretreatment of in vitro translated Plsp1 with buffer, SE, or Cpn60_14_ occurred for 0, 10, 30, or 60 minutes (Fig. 7A-B). After 60 minutes, only 68% of buffer-pretreated Plsp1 remained transport competent (Fig. 7B). Incubation with either SE or Cpn60_14_ did not provide detectable enhancement to transport competence at any time point tested (Fig. 7A-B). While Plsp1 interacts with Cpn60 in the stroma, this interaction cannot protect Plsp1 from losing its ability to integrate over time.

## Discussion

Approximately 90% of thylakoid TM proteins transit the aqueous stroma prior to integration, assisted by largely unidentified mechanisms. The nuclear-encoded Plsp1 binds to Cpn60, as demonstrated by in vitro pulldown, reconstitution, and Proteinase K protection assays (Fig. 1). Plsp1 pulls down the abundant Cpn60 α1 and β1 isoforms, but not the substrate-specialized α2 or β4 (Peng et al., 2011; Ke et al., 2017), indicating Plsp1 is a substrate of the major “housekeeping” Cpn60 oligomer. Chaperonins bind and encapsulate proteins, typically leading to their substrate’s assumption of a native conformation, though this often requires multiple cycles of encapsulation (Hayer-Hartl et al., 2016). The increased integration resulting from RbcL displacing Plsp1 from Cpn60 (Fig. 6D-E) suggests that Cpn60-bound Plsp1 retains a non-native confirmation. Cpn60 interacts with the Plsp1 sequences which will fold to form its hydrophobic membrane association surface (Fig. 3B-C). Since lipid association is critical for folding catalytically active Plsp1 (McKinnon et al., 2020), it may be unfavorable for Plsp1 to fold within the chaperonin. Unfolded Plsp1 bound by a Cpn60 α1β1 oligomer would remain competent for thylakoid insertion.

Our in vitro targeting assays in isolated thylakoids (Fig. 5-6) and intact chloroplasts (Fig. 2) support Cpn60 holding Plsp1 transiently prior to its thylakoid insertion. Three models could explain the hand-off of Plsp1 from Cpn60 to cpSec1. First, Cpn60 could bind to cpSec1 components and pass Plsp1 to the translocon. In this model, Cpn60 functions analogously to cpSRP or SecB, the *E. coli* targeting factor which delivers unfolded secreted proteins but which is not conserved in chloroplasts (Dempsey et al., 2004; Sala et al., 2014). We did not detect interaction between cpSecA1 and Cpn60 in the stroma in the absence of Plsp1 (Supplemental Fig. 6), though we cannot exclude Plsp1-mediated interaction between these proteins. If Cpn60 behaved like cpSRP or SecB, we would expect Cpn60 to stimulate integration of Plsp1, which is inconsistent with our in vitro transport assay results (Fig. 5). Second, Cpn60 could pass Plsp1 to another stromal factor in order for this additional component to deliver Plsp1 to cpSec1. The known additional stromal protein involved in targeting Plsp1 is cpSecA1. However, while cpSecA1 interacts with its substrates at the thylakoid membrane, cpSecA1 has not been detected binding its substrates in the stroma to recruit them to the membrane (Haward et al., 1997; Ma, 2000). We did not identify another stromal protein in our pulldown (Fig. 1) or complex in our import-chase assays (Fig. 2) that appeared to act as an intermediate between Cpn60 and cpSec1. These two models are not supported well by our data.

The third hand-off mechanism is a catch-and-release model in which Plsp1 cycles on and off Cpn60, rebinding Cpn60 until encountering cpSec1. During in vitro transport in isolated thylakoids, Cpn60 reduced Plsp1 integration (Fig. 5). ATP hydrolysis and displacement by the obligate substrate RbcL released Plsp1 from Cpn60 for insertion into the thylakoid (Fig. 6). The effect of RbcL can be attributed to two processes, which are not mutually exclusive. First, occupying Cpn60 with RbcL lowers the apparent concentration of free Cpn60 to bind Plsp1. Second, given that chaperonins preferentially bind obligate substrates like RbcL over less stringent substrates (Kerner et al., 2005), RbcL will out-compete Plsp1 and more readily rebind Cpn60 during chaperonin cycling. Increasing cpSecA1 also partially restored integration (Fig. 5E-F), suggesting that Plsp1 exists in an equilibrium between Cpn60- and cpSecA1-bound forms. The correlation between reduced Cpn60 interaction and increased dp1 during import-chase experiments (Fig. 2C) implies that, ultimately, cpSecA1 collects the imported Plsp1 recently released from Cpn60. Our data best support the catch-and-release model in which Cpn60 captures and releases Plsp1 cyclically, while cpSec1 recognizes free stromal Plsp1 and integrates it. The cpSec1 translocon depletes the free Plsp1 from the stroma, and by mass action, Cpn60 continuously releases its Plsp1 population until integration is complete.

Involvement of Cpn60 in cpSec1-mediated targeting was unexpected, considering its classical role as a folding chaperone in cooperation with Cpn10/20 and ATP. Most known Cpn60 substrates are soluble stromal proteins. Consistent with Cpn60’s folding function, biochemical data demonstrates Cpn60 binds and likely assists folding of phytoene desaturase (PDS) and the Rieske iron-sulfur protein (PetC), and this interaction promotes membrane association for PDS and chloroplast twin-arginine translocon-mediated thylakoid insertion for PetC (Madueno et al., 1993; Bonk et al., 1997, Molik et al., 2001). Chaperonins can, in certain cases, function in membrane protein insertion in vitro by stabilizing exposed hydrophobic residues. GroEL maintains the solubility of multispan TM proteins bacteriorhodopsin (BR) and *E. coli* lactose permease (LacY) and delivers both of these proteins to membranes (Bochkareva et al., 1996; Deaton et al., 2004). Neither Plsp1’s primary nor tertiary structure resemble BR or LacY, as Plsp1 has a single TMD and a large lumenal domain. The transfer of BR from GroEL to inverted membrane vesicles occurred without addition of other factors but was enhanced by GroES and ATP (Deaton et al., 2004). LacY’s transfer was also enhanced by GroES, but in this case, ATP was strictly required (Bochkareva et al., 1996). Similarly, the transfer of Plsp1 from Cpn60 to thylakoids was inefficient in the presence of Cpn20 and ATP (Fig. 5).

Efficient hand-off of Plsp1 to cpSec1 was influenced by displacement of Plsp1 from Cpn60 by other chaperonin substrates. Our in vitro binding and transport assays used RbcL because it is firmly established as an obligate Cpn60 substrate. In our second-dimension analysis, we observed RbcL corresponding to approximately 20% of the assembled RbcL pool co-migrating with Cpn60 (Supplemental Fig. 3B), consistent with other reports (Peltier et al., 2006; Olinares et al., 2010; Feiz et al., 2012; Feiz et al., 2014), and RbcL was abundant in our excised 700-kDa Cpn60 complex (Fig. 2). We also detected minor levels of other known Cpn60 substrates co-migrating with the 700-kDa complex: NdhH (Peng et al., 2011), CF1-α, CF1-β (Chen and Jagendorf, 1994; Mao et al., 2015), and glutamine synthetase (Lubben et al., 1989) (Supplementary Data Set 2). In *E. coli*, experiments suggest that 99% of available chaperonins are occupied by substrates (Kerner et al., 2005; Jewett and Shea, 2010). Our data are likewise consistent with a heavily occupied Cpn60.

For RbcL, its physiological concentrations are not dissimilar to the molar excess of chemically denatured RbcL we added in vitro. The steady-state concentration of active RbcL in chloroplasts was measured at approximately 4 mM (Jensen and Bahr, 1977). In slow-growing *Arabidopsis* leaves, 3% of the RbcL pool was replaced per day; in rapidly-growing *Arabidopsis* leaves, the fraction of newly synthesized RbcL increased to 35% per day (Li et al., 2017). Assuming 13 μM Cpn60 in the chloroplast (Musgrove et al., 1987), each Cpn60 must fold 0.4 to 5 RbcL per hour. Estimated rates of RbcL folding by chaperonins (approximately 0.2 min^-1^) (Viitanen et al., 1995; Liu et al., 2010; Hauser et al., 2015) suggest that newly synthesized RbcL will occupy 3-40% of a single Cpn60’s capacity per hour, depending on growth rates. The targeting of Plsp1 is likely highest in growing leaves (Schmid et al., 2005; Hsu et al., 2011) that also have high RbcL synthesis rates (Lorimer, 1996; Li et al., 2017). Additionally, it has been clear from the first report of RbcL interaction with Cpn60 (Barraclough and Ellis, 1980) that Rubisco holoenzyme assembly, not RbcL synthesis, is rate-limiting. Rubisco assembly requires Cpn60 to fold RbcL, then four other chaperones to construct the RbcL_8_-RbcS_8_ complex, and the *in vivo* kinetics of this process remain opaque (Vitlin Gruber and Feiz, 2018). If the assembly chaperones are unavailable, RbcL remains bound to Cpn60 (Feiz et al., 2012; Feiz et al., 2014). There is clearly enough Cpn60 free to interact with Plsp1 in the stroma, but sufficiently high concentrations of RbcL and other endogenous substrates are present to displace Plsp1 from Cpn60.

Biochemical and genetic evidence suggests a limited role for GroEL in stabilizing transport-competent states of secreted proteins. The periplasmic β-lactamase and outer membrane OmpA, both exported by Sec in *E. coli* (Brundage et al.; Pradel et al., 2009), were identified as *in vivo* GroEL substrates, along with 13 other periplasmic and outer membrane proteins (Kerner et al., 2005). Pretreatment with GroEL maintains the transport-competence of pre-β-lactamase (Bochkareva et al., 1988) and proOmpA (Lecker et al., 1989). Conditional GroEL/ES mutants exhibit delayed pre-β-lactamase secretion (Kusukawa et al., 1989), and GroEL overexpression restored β-lactamase export (Kusukawa et al., 1989), improved secretion of export-deficient LamB-LacZ fusions (Phillips and Silhavy, 1990), and partially complemented conditional SecA, SecY, and SecE mutants (Van Dyk et al., 1989; Danese et al., 1995; Castanié-Cornet et al., 2014).

Does Cpn60 serve as a simple waystation or does it actively maintain Plsp1’s transport competence? Our data present a complex picture. Where cpSRP43 can prevent LHCP aggregation and rescue aggregated LHCP, maintaining complete transport competence over 1 hour of room temperature pretreatment, we were unable to detect any stromal protein, including Cpn60, that performed this function for Plsp1 (Fig. 7). If Plsp1 remains bound to Cpn60 but loses its insertion competence, Cpn60 may be able to reroute Plsp1 to a quality control pathway through its interaction with FtsH11 (Adam et al., 2019). Maintenance of competence by Cpn60 may be detected under more stressful in vitro conditions (for example, higher temperature), although in chloroplasts experiencing such stress, Cpn60’s clientele would expand globally, reducing its ability to protect Plsp1 in particular. However, when Plsp1 loses the ability to interact with Cpn60 through its hydrophobic membrane association surface (β1-2) or with cpSec1 through its thylakoid targeting signal (TMD), its thylakoid integration is abolished (Fig. 4). Δβ1-2 Plsp1 fractionates with membranes (Fig. 4B), and its failure to integrate suggests that it assumes a transport-incompetent conformation, although it is also possible that this deletion compromises the targeting signal. Cpn60 interaction is not necessary for Plsp1 to integrate in vitro, nor does it extend Plsp1’s transport competence in vitro, but in intact chloroplasts, losing Cpn60 interaction correlates with compromised integration.

Cpn60 is essential, which precluded the use of genetic approaches to study the importance of Cpn60 for Plsp1’s targeting. Cpn60 α1 mutants exhibit embryo lethality in *Arabidopsis* and seedling lethality in maize (Barkan, 1993; Feiz et al., 2012) and rice (Kim et al., 2013). Double mutants of redundant β1 isoforms in *Arabidopsis* result in seedling lethality (Suzuki et al., 2009; Ke et al., 2017). Mild Cpn60 β1 knockdown in tobacco causes leaf chlorosis and starch accumulation in chloroplasts, suggesting that chloroplast development is perturbed (Zabaleta et al., 1994). In principle, inducible Cpn60 knockdown mutants could be generated to examine the effect on Plsp1’s targeting and on thylakoid development in general, but such study is outside the scope of this work. Mutants with impaired chloroplast proteostasis, including those disrupting the cpSRP pathway, often show increased accumulation of Cpn60 and other chaperones to compensate (Rutschow et al., 2008; Nishimura and van Wijk, 2015). While Cpn60 is not the only chaperone available to assist thylakoid TM protein targeting, Cpn60 provides a possible path for proteins transiting the stroma.

In summary, our results demonstrate interaction between Plsp1 and Cpn60 and support a mechanism by which Cpn60 holds Plsp1 until the cpSec1 translocon recruits it to the membrane for integration.

## Methods

### DNA constructs

Plasmids encoding prPlsp1 and mPlsp1 for in vitro translation (Endow et al., 2015) and mOE33 (Chou et al., 2006) and cpSecA1 (Endow et al., 2015) for recombinant expression in *E. coli* were described previously. To generate the internal prPlsp1deletions ΔTMD (residues 112-131) and Δβ1-2 (132-157), two fragments were amplified from prPlsp1 in pIVTGW-SP6 with the indicated primers (Supplementary Table 1), then combined by long-flanking homology PCR (Tian et al., 2004; Endow et al., 2015). To generate C-terminal prPlsp1 deletions ΔC (272-291) and Δβ9-C (262-291), prPlsp1 was amplified using the indicated primers complementary to the new 5’ end. Plsp1 deletion constructs were subcloned into pIVTGW-SP6 by Gateway cloning for in vitro translation. mPlsp1_ΔTMD_ was amplified from prPlsp1_ΔTMD_ in pIVTGW-SP6 with primers containing NdeI and BamHI restriction sites, digested, and ligated into similarly digested pET16b. Untagged *P. sativum* Cpn60α1 (Psat7g144320) and Cpn60β1 (Psat1g001680) constructs were a gift from A. Azem (Dickson et al., 2000). To add purification tags, Cpn60α1 in pET24a was excised with NdeI and BamHI, then ligated into similarly digested pET16b. Cpn60β1 was amplified from pET24d with primers adding a 5’-NdeI and 3’-XhoI site and cloned into pCR2.1 using an Invitrogen TOPO-TA kit. Cpn60β1 was digested with NdeI and XhoI, then ligated into similarly digested pET16b.

### Protein preparation

For radiolabeled proteins, constructs were translated in vitro using a TNT (TM) Coupled Rabbit Reticulocyte Lysate kit (Promega). His-mPlsp1_ΔTMD_, His-mOE33 (Chou et al., 2006), His-Cpn60 α1, His-Cpn60 β1, and His-cpSecA1 (Endow et al., 2015) were expressed in *E. coli* and purified by Ni-NTA affinity chromatography. Full expression and purification details may be found in Supplemental Materials. Protein purity was assessed by SDS-PAGE and concentration by Bradford assay (Bio-Rad).

### Isolation of chloroplasts, thylakoids, and concentrated stromal extract

Intact chloroplasts were isolated from 9 to 13-day-old *P. sativum* seedlings as described (Inoue and Keegstra, 2003). Preparation of thylakoids and concentrated stromal extract (SE, at 2.5 mg chlorophyll equivalent chloroplasts/ml) was as described (Endow et al., 2015; Asher et al., 2018).

### In vitro pulldown assay

In vitro pulldown was modified from (Flores-Pérez et al., 2016) and (Peng et al., 2011). 100 μl SE was mixed with 20 μg of recombinant protein in binding buffer: 50 mM HEPES pH 8.0, 3.3 mM Tris, 330 mM sorbitol, 20 mM imidazole, 13.3 mM NaCl, 10 mM MgCl_2_, 10 mM ADP, 20 mM glucose, and 20U/ml hexokinase (Roche). The sample was incubated with rocking at 4°C for 1 hour, then added to 20 μl Ni-NTA equilibrated with binding buffer without ADP, glucose, or hexokinase. Proteins were bound to Ni-NTA overnight at 4°C with rotation. The solution was transferred to a 200 μl column (Promega). Unbound proteins were collected by centrifugation at 800 *×g* for 1 min, then resin was washed with 3 ml of SE buffer: 50 mM HEPES pH 8.0, 330 mM sorbitol, 10 mM MgCl_2-,_ and 20 mM imidazole. For sequential elution, resin was incubated with 60 μl of 10 mM Mg-ATP or 60 μl 300 mM imidazole in 50 mM HEPES pH 8.0, 330 mM sorbitol, 10 mM MgCl_2_ for five minutes before centrifugation. Aliquots were analyzed by SDS-PAGE and immunoblotting. For LC-MS/MS analysis, 50% of the first imidazole elution was run into the resolving gel on SDS-PAGE, stained, and excised.

### In vitro chaperonin oligomer reconstitution assay

In vitro reconstitution was performed as in (Dickson et al., 2000; Vitlin et al., 2011; Gruber et al., 2014): 50 μM Cpn60 α1 and β1 and 25 μM Cpn20 were incubated in 50 mM Tris-HCl pH 8.0, 300 mM NaCl, 10 mM MgCl_2_, 4 mM KCl, 2 mM DTT, and 1.6 mM MgATP at room temperature for 5 min, then at 30°C for 1 hour. To separate tetradecameric (Cpn60_14_) Cpn60 from other oligomers, the reconstitution reaction was separated by size-exclusion chromatography on a GE Superose 6 column in 50 mM HEPES-KOH pH 8.0, 100 mM NaCl, 5% glycerol (HVLC) at room temperature. The oligomeric state was confirmed by separating 5 ul equivalents of fractions on 6% native-PAGE. Fractions containing Cpn60_14_ were pooled and concentrated in an Amicon 30K MWCO filter. Concentrated Cpn60_14_ was quantified by Bradford assay (Bio-Rad). The calculated molecular weights of His-tagged Cpn60α and β monomers were used to calculate the molecular weight of an α7β7 oligomer, and concentrations are presented in terms of oligomer unless otherwise stated.

### In vitro chaperonin binding

Chaperonin binding assay reactions included 200 nM Cpn60_14_ and 5% radiolabeled mPlsp1 in 50 mM HEPES-KOH, pH 8.0, 330 mM sorbitol (IB). For experiments in Fig. 1B, reactions were incubated for 30 min at room temperature. Addition of 10 mM MgATP or AMP-PNP was followed by incubation for 10 min at room temperature. 5 μl of binding reaction was separated by 4-14% BN-PAGE, as described (Endow and Inoue, 2013).

### Proteinase K protection time course

Proteinase K protection time courses were performed at room temperature as in (Mao et al., 2015) with modifications. Treatments were (a) Cpn60_14_, (b) Cpn60_14_ and Cpn20, (c) Cpn20, (d) BSA and Cpn20, or (e) buffer only (no addition). Radiolabeled Plsp1, produced by in vitro translation, was incubated with Cpn60_14_, BSA, or buffer for 30 min. ADP was added to all treatments, and Cpn20 was added to designated treatments, followed by a 10 min incubation. The protease protection assay was initiated by the addition of 1.2 ug/ml Proteinase K (Sigma P2308). Aliquots were removed at 0, 10, 20, and 60 minutes and immediately quenched by addition of 1 mM PMSF in isopropanol. Final concentrations of components were 2% substrate, 0.8 μM Cpn60_14_, 2 μM Cpn20, 11.2 μM BSA (equivalent to Cpn60_14_ in terms of monomer), and 1 mM ADP in ProtK buffer (50 mM Tris-HCl pH 7.5, 50 mM KCl, 5 mM MgCl_2_).

The samples were separated on 12% SDS-PAGE alongside 100% and 50% substrate input standards. Radioactive signals were quantified by densitometry in ImageJ. The protected fraction was calculated for each treatment, normalized to the mean of the 0-min time points per treatment. Data were fit to a falling single exponential in Kaleidograph (version 4.1.1).

### In vitro protein import

Protein import assays with intact chloroplasts were performed as described (Inoue and Keegstra, 2003). Briefly, reactions containing chloroplasts at 0.25 mg/ml chlorophyll (Chl), 3 mM MgATP, and 10% translation in import buffer (IB, 50 mM HEPES-KOH pH 8.0, 330 mM sorbitol) were incubated in 20-40 μE/m^2^s light. Intact chloroplasts were reisolated through 40% Percoll in IB. An aliquot of each import was used to quantify chloroplast recovery; the remainder was pelleted at 2,000 ×*g* and either used for subsequent analysis or resuspended to 0.3 mg/ml Chl in SDS-PAGE sample buffer.

The import-chase assay was as described with minor modifications (Endow et al., 2015). 1 mM lincomycin was included throughout import and chase in a subset of experiments to suppress chloroplast translation (Kim et al., 1994; Chotewutmontri and Barkan, 2018). After a 10-min import assay, chloroplasts were diluted in ice-cold IB to less than or equal to 0.5 mM ATP, pelleted, and treated with 0.2 μg/μl thermolysin to remove unincorporated precursor. Thermolysin activity was quenched with 10 mM EDTA, and intact chloroplasts were recovered through 40% Percoll, washed, and divided into three equal aliquots. 3 mM ATP addition initiated the chase, with chase incubations in light staggered to end simultaneously. An aliquot (approximately 4 μg Chl) of intact chloroplasts from each chase was reserved as total import (T), and the remainder was lysed in 10 mM HEPES-KOH pH 8.0, 10 mM MgCl2 (HM) at 1 mg/ml initial Chl for 10 min on ice. The soluble fraction was collected by centrifugation at 16,000 *×g* for 20 min and clarified of any residual membranes in a second 16,000g, 20 min centrifugation. The pellet was treated with 0.1 μg/ul thermolysin for 40 min on ice. For the experiments quantified in Fig. 2C, the supernatant was adjusted so that 3 μg Chl was analyzed by 4-14% BN-PAGE. To load on BN-PAGE, supernatant was mixed with two volumes of BN-PAGE loading solution.

Fractionation into soluble (S1), alkaline-extracted (S2), and alkaline-resistant (P) samples was as described (Inoue et al., 2006). Briefly, chloroplasts were lysed in HM and the soluble (S1) fraction was collected by centrifugation at 16,000 *×g* for 20 min. The pellet was washed in 0.1 M Na_2_CO_3_ for 10 min on ice, and the alkaline-extracted (S2) fraction collected by centrifugation at 16,000 *×g* for 20 min. The pellet (P) represented alkaline-resistant proteins. When soluble complexes were analyzed by BN-PAGE (Fig. 3), reisolated chloroplasts were lysed in HM at 1 mg/ml initial Chl concentration. As described for import-chase, the soluble fraction after centrifugation was spun for a further 20 min at 16,000 *×g* to pellet any residual membranes.

### In vitro transport

Protein transport assays with isolated thylakoids were performed as described (Endow et al., 2015; Asher et al., 2018). In brief, reactions contained thylakoids at 0.33 mg/ml Chl, 5 mM MgATP, and 10% translation product and were incubated at room temperature for 30 min in 70-100 μE/m^2^s light. Integration was assessed by treatment with 0.1 μg/μl thermolysin for 40 min on ice. After thermolysin activity was quenched with EDTA, recovered membrane pellets were analyzed by SDS-PAGE. When cpSecA1, Cpn60 Cpn60_14_, or Rubisco were included, control reactions were supplemented with their respective buffers.

### Rubisco displacement assays

Spinach Rubisco (Sigma R8000, RbcL_8_:Rbcs_8_) was denatured in 6M guanidine-HCl, 5 mM DTT for 30 min at room temperature. RbcL and RbcS were quantified by densitometry relative to BSA, and the calculated concentration of RbcL was reported. Whenever RbcL was added, an equal concentration of denatured RbcS was also present. The final concentration of guanidine-HCl in binding and transport assays was 70 mM, which did not inhibit Cpn60-Plsp1 binding and inhibited transport by 20%. To minimize guanidine-HCl addition, denatured RbcL was diluted to 2.5 M guanidine-HCl immediately before its addition to binding or transport assays. 6 μM RbcL was the highest final concentration which could be added without immediate aggregation.

For RbcL displacement analysis in chaperonin binding assay, conditions were modified from (Liu et al., 2010). Reactions contained 200 nM Cpn60 Cpn60_14_, 5 mM DTT, 1 mg/ml BSA, and 5% radiolabeled mPlsp1. Reactions were incubated at room temperature for 5 min, then chilled on ice for 5 min. Denatured RbcL was diluted 36-fold into the chilled reaction, which was transferred back too room temperature for 20 min. 5 μl of binding reaction was analyzed by 4-14% BN-PAGE. Each independent experiment was performed in triplicate for each RbcL concentration and in duplicate for mock buffer addition.

To analyze RbcL’s effect on in vitro transport, denatured RbcL was diluted 36-fold to 3 μM at t = 0 or t = 10 min of a 30-min transport assay. When RbcL was added at t = 0 min, it was the last component added to an ice-cold transport reaction before the reaction was transferred to light at room temperature; when RbcL was added at t = 10 min, the transport reaction was equilibrated to room temperature.

### LC-MS/MS and data analysis

In-gel trypsin digestion and LC-MS/MS were performed at the UC Davis Proteomics Core Facility. Peptide identification and label-free quantification (LFQ ratio equal to 1) were performed in MaxQuant (1.5.6.0) using default parameters (Cox and Mann, 2008). Full details describing generation of a *P. sativum* search database and data processing and analysis may be found in the Supplemental Methods.

### Stromal concentration of Cpn60_14_ and cpSecA1 estimation

The crude approximation of the stromal concentration of Cpn60_14_ and cpSecA1 were based off a method used to approximate RbcL concentrations (Jensen and Bahr, 1977). The estimated μl stroma per mg Chl volume used were 20, 25, or 40 μl per mg Chl (Heldt, 1980; Howitz and McCarty, 1982; Robinson, 1984). The molecular weight of mature cpSecA1 and Cpn60α1 was predicted by Expasy. The Cpn60_14_ oligomer calculation assumed that the stromal oligomer contains 6 Cpn60α1 protomers, consistent with our data (Supplemental Data Set 2) and LC-MS/MS-based approximations in *Chlamydomonas* (Bai et al., 2015; Zhao et al., 2019).

### Antibodies

Antibodies used in this work were anti-Hsp60 (Enzo), anti-His (Santa Cruz), anti-OE33 (Chou et al., 2006), anti-cpSecA (Nakai et al., 1994), and anti-Cpn60α1 (Agrisera).

### Nomenclature

In this study, we refer to Cpn60 isoforms according to the convention used by TAIR for *Arabidopsis thaliana* that orders isoforms by expression level. Supplementary Table 2 lists the *Pisum sativum* gene identifiers (v1a, (Kreplak et al., 2019)), *P. sativum* UniProt identifiers, and corresponding *Arabidopsis thaliana* gene identifiers (Araport 11) for each Cpn60 isoform.

## Supplemental Data

Supplemental Methods: Expanded procedures for protein purification and LC-MS/MS analysis

Supplemental Data Set 1: Full list of proteins identified in mPlsp1_ΔTMD_ in vitro pulldown.

Supplemental Data Set 2: Full list of proteins identified in excised 550- and 700-kDa complexes

Supplemental Figure 1: mPlsp1 is protected from Proteinase K by Cpn60/Cpn20

Supplemental Figure 2: The 180-kDa and 550-kDa complexes during the import-chase assay

Supplemental Figure 3: Lincomycin suppresses the appearance of the 550-kDa complex in an import-chase assay

Supplemental Figure 4: The 550-kDa and 180-kDa bands do not require any of the Plsp1 sequences analyzed.

Supplemental Figure 5: Δβ1-2 remains thermolysin-susceptible after 30-minute import

Supplemental Figure 6: cpSecA1 does not pull down Cpn60 from stromal extract

Supplemental Figure 7: Stromal concentrations of cpSecA1 and Cpn60 are high

Supplemental Figure 8: High concentrations of protein do not inhibit integration of Plsp1.

Supplemental Figure 9: RbcL displacement is less effective at room temperature.

Supplemental Table 1: Cloning primers

Supplemental Table 2: Cpn60 gene identifiers in *Pisum sativum* and *Arabidopsis thaliana*

### Acknowledgements

We gratefully acknowledge Joshua K. Endow for his critical feedback and advice on many experiments. A. Azem kindly provided a gift of the Cpn60 α1 and β1 constructs. Brett Phinney and Michelle Salami at the UC Davis Proteomics core provided training, technical expertise, and assistance with sample preparation and analysis for LC-MS/MS experiments. Prakitchai Chotewutmontri provided advice on the use of lincomycin, and Manajit Hayer-Hartl, and F. Ulrich Hartl gave critical feedback on chaperonin experiments, particularly the Rubisco displacement assay. We also thank Philipp Zerbe and Rebecca Roston for their critical reading of this manuscript. This work was supported by the Office of Basic Energy Sciences of the U.S. Department of Energy (grant DE-SC0017035 to S.M.T. and grant DE-FG02-08ER15963 to K.I.), the UC Davis Department of Plant Sciences (Graduate Student Research Assistantship and Henry A. Jastro Graduate Research Award to L.K.), and the UC Davis Department of Plant Biology (Stocking Fellowship to L.K.).

This is the last of three research papers that were published in the past year listing as co-author our friend, mentor, and colleague Kentaro Inoue, who was tragically killed in an accident three years ago. These papers were each born of his inspiration and ideas, and the first author of each was one of the three graduate students in his lab at the time of his death. To mark the end of this chapter of Kentaro’s active contributions to science, we would like to dedicate this paper to his memory.

## Author Contributions

L.K, K.I, and S.M.T. designed the experiments. L.K. performed all experiments and analyzed the data, and L.K. and S.M.T. wrote the manuscript.

## Figure Legends

**Supplemental Data Set 1**: Full list of proteins identified in mPlsp1_ΔTMD_ in vitro pulldown. Proteins identified in LC-MS/MS analysis of proteins co-eluting with Plsp1. 50% of the first imidazole elution for the pulldown (mPlsp1_ΔTMD_ bait) and control (SE-only mock) from two independent experiments was analyzed. Curated_Name was assigned from BLASTX annotations from Swiss-Prot or Trembl for the Cool Season Food Legume (CSFL) *Pisum sativum* reference transcriptomes and manual BLAST of *P. sativum* sequences against *Arabidopsis thaliana* proteins. Psat_ID indicates the gene model from the *P*.*sativum* genome (v1a); “unidentified” indicates proteins identified from transcriptomic sequences only. The average Fold_Change between pulldown (mPlsp1_ΔTMD_ bait) and control (SE-only mock) was calculated from the log2 transformed ratio of iBAQ values for each protein; “IP_only” indicates detection solely in the pulldown. Enriched (+) indicates proteins which were significantly more abundant in the pulldown than the control; enrichment was determined by Student’s t-test (p < 0.05) with correction for multiple comparisons. The % relative to total Cpn60 (%_Rel_Total_Cpn60) was calculated using the total iBAQ value for all detected Cpn60 isoforms. Peptides indicate the number of peptides matched to a protein, while Unique_peptides are the number of matched peptides unique to that protein. Sequence_coverage indicates the percentage of a protein’s sequence identified by matched peptides. Protein_IDs are all *P. sativum* genes and translated CSFL transcripts conglomerated in MaxQuant for a particular protein’s identification. The log_2_ transformed and raw iBAQ values for both pulldown (IP) and control (Ctrl) replicates are included.

**Supplemental Data Set 2**: Full list of proteins identified in excised 550- and 700-kDa complexes.

Proteins identified in LC-MS/MS analysis of 550-kDa and 700-kDa regions. The 700-kDa band was excised from the soluble fraction after 10 min import of non-radiolabeled Plsp1; the 550-kDa band was excised from the soluble fraction after 10 min import and 30 min chase of non-radiolabeled Plsp1. Data represent two independent experiments for each band, and independent experiments were performed in triplicate (700-kDa) or duplicate (550-kDa). Curated_Name is the name of the protein. Curated names, Psat_ID, Peptides, Unique_peptides, Sequence_coverage, and Protein_IDs were as described for Supplemental Data Set 1. For Cpn60 isoforms, *P. sativum* gene names are included in parentheses, and “unidentified” indicates Cpn60s identified from transcriptomic sequences only. 700_unadjusted and 550_adjusted represent the mean log_2_ iBAQ intensities for a protein. 700_Rnorm and 550_Rnorm represent those mean log_2_ iBAQ intensities relative to the mean intensity of RbcL in the 550-kDa band, which was the most abundant protein in either band. The unadjusted log2 iBAQ values for each band and replicate are included in columns that follow the naming convention: BAND_ EXPERIMENT.REPLICATE.

**Supplemental Figure 1**: mPlsp1 is protected from Proteinase K by Cpn60/Cpn20. Proteinase K protection assay of Cpn60-mediated substrate encapsulation. mPlsp1 was incubated with Cpn60 tetradecamer (0.8 μM), BSA (11.2 μM, equivalent to Cpn60 protomer concentration), or buffer for 30 min at room temperature to allow Plsp1 to bind Cpn60. ADP and, where indicated, Cpn20_4_ (2 μM), were added and incubated for 10 min. Assay was initiated with the addition of 1.2 ug/ml Proteinase K, and aliquots were removed at 0, 10, 20, and 60 minutes. After separation on SDS-PAGE, aliquots were visualized by autoradiography. Protection at each time point for each treatment was quantified relative to the 0-minute time point. Points represent mean; error bars represent ± 1 SD. Lines are drawn to aid visualization. N = 3 independent digests.

**Supplemental Figure 2**: The 180-kDa and 550-kDa complexes during the import-chase assay Quantification of 550- and 180-kDa complexes. Import-chase assays were performed without lincomycin and separated 3 μg chlorophyll on BN-PAGE. The occupancy of the 550- and 180-kDa complexes at each chase time point was quantified relative to t=0 min. N = 10.

**Supplemental Figure 3**: Lincomycin suppresses the appearance of the 550-kDa complex in an import-chase assay

(A) Import-chase assay followed by BN-PAGE analysis. Radiolabeled Plsp1 was imported into illuminated (50 μE/m^2^s) chloroplasts with 3 mM ATP for 10 min, followed by treatment with thermolysin for 30 min on ice in the dark to remove unincorporated precursor. The protease was quenched, and intact chloroplasts were reisolated and incubated in 50 μE/m^2^s light with 3 mM ATP for 0, 10, or 30 min. One import-chase assay included 1 mM lincomycin throughout (lanes 1-3); the second import-chase assay did not include lincomycin (lanes 4-6).

(B) 2D analysis of import-chase BN-PAGE. Duplicate lanes from the import-chase assay shown in (A) were run on 2D SDS-PAGE. The Coomassie-stained gel for t=0 min (top) and the phosphorimages of all three chase times are shown. Regions corresponding to the 700-kDa complex, 550-kDa complex, and dye front are indicated with blue underlay. Sizes of Cpn60, RbcL, and Plsp1 (pr, precursor; m, mature) are indicated. Orange circles on 30-min chase indicate chloroplast-synthesized RbcL migrating as its assembled holoenzyme.

**Supplemental Figure 4**: The formation of the 550-kDa and 180-kDa bands do not require any of the Plsp1 sequences analyzed.

To quantify the 550-kDa complex (A) and 180-kDa complex (B), constructs were analyzed pairwise with their full length Plsp1 control. Densitometry signals for each deletion construct and the full length Plsp1 control were normalized to account for different numbers of methionine residues and import efficiency. Signals from an individual experiment were normalized to the mean of both signals from that experiment. Normalized signals for full length Plsp1 and a deletion construct were then divided by the mean of signals for full length Plsp1 for all replicates. Significant differences (p<0.05, *) between mean full length and deletion construct were calculated by t-test.

**Supplemental Figure 5**: Δβ1-2 remains thermolysin-susceptible after 30-minute import.

(A) Thermolysin resistance of prPlsp1 after extended in vitro import. In vitro-translated, radiolabeled Plsp1 was imported into illuminated (50 μE/m^2^s) chloroplasts with 3 mM ATP for 30 min (left) or 10 min (right). Chloroplasts were pelleted (Tot) or lysed and treated with thermolysin (Tm). Equal chlorophyll (Chl) equivalents were run on SDS-PAGE alongside 10% input translation and visualized by autoradiography. Precursor (pr), mature (m), dp1, and dp2 are indicated.

(B) Thermolysin resistance of Δβ1-2 after extended in vitro import. In vitro-translated, radiolabeled Δβ1-2 was imported into illuminated (50 μE/m^2^s) chloroplasts with 3 mM ATP for 10 min or 30 min. Chloroplasts were pelleted (Tot) or lysed and treated with thermolysin (Tm) and analyzed as in (A). Precursor (pr), mature (m), and potential dp1 and dp2 are indicated. Black lines mark removed lanes. Experiment performed in triplicate.

**Supplemental Figure 6**: cpSecA1 does not pull down Cpn60 from stromal extract.

In vitro pulldown of Cpn60 from stromal extract (SE) with His-tagged bait. 20 μg cpSecA1 was incubated with SE equivalent to 250 ug Chl, then bound to Ni-NTA, washed, and sequentially treated with 10 mM MgATP and 300 mM imidazole. Representative immunoblots of two independent experiments show SE input (1%), bait input (2.5%), unbound (1%), wash (25%), ATP treatment (ATP, 25%), and imidazole elution (elute, 25%). Cpn60 was detected with anti-Hsp60; His-cpSecA1 was detected with anti-His.

**Supplemental Figure 7**: Stromal concentrations of cpSecA1 and Cpn60 are high.

(A) Semi-quantitative cpSecA1 immunoblot of five SE samples from independent chloroplast isolations. Standards are 100 and 50 ng recombinant cpSecA1, which migrates at a larger size because of its purification tags. Positions of empty lanes removed to condense the image are indicated with black lines.

(B) Semi-quantitative Cpn60α1 immunoblot of five SE samples from independent chloroplast isolations. Standards are 0.0625, 0.125, 0.25, 0.5, and 1 μg recombinant Cpn60α1, migrating at a larger size because of its His tag. Positions of lanes removed to condense the image are indicated with black lines.

(C) Approximation of the Cpn60_14_ and cpSecA1 concentration in the chloroplast, based on estimated stromal volumes of 20, 25, or 40 per mg Chl (Heldt, 1980; Howitz and McCarty, 1982; Robinson, 1984).

**Supplemental Figure 8**: High concentrations of protein do not inhibit integration of Plsp1. In vitro transport assay with BSA equimolar to 560 nM Cpn60_14_. Radiolabeled Plsp1 was incubated with thylakoids in the presence of 5 mM ATP (lanes 2-3), 200 nM cpSecA1 (lanes 2-3), and 7.84 μM BSA (lane 3) for 30 min in 90 μE/m^2^s light. Thylakoids were treated with thermolysin and analyzed by SDS-PAGE and autoradiography alongside 5% input translation (tl). Sizes of mature Plsp1 (mPlsp1), dp1, and dp2 are indicated. Representative phosphorimage of two independent experiments, performed in triplicate.

**Supplemental Figure 9**: RbcL displacement is less effective at room temperature.

Competition binding assay. In vitro-translated, radiolabeled Plsp1 was incubated with 200 nM Cpn60_14_ for 10 min at room temperature, then 0, 0.15, 1.5, or 3 μM guanidine-denatured RbcL (lanes 1-4, respectively) was diluted 36-fold into the reaction at room temperature. Reactions were incubated for 20 minutes further at room temperature, then analyzed by BN-PAGE and autoradiography. Representative phosphorimage of 2 independent experiments; exact RbcL concentrations differed between independent experiments.

**Supplemental Table 1**: Cloning primers.

Forward and reverse primers used in the preparation of constructs in this work.

**Supplemental Table 2**: Cpn60 gene identifiers in *Pisum sativum* and *Arabidopsis thaliana*. For each Cpn60 isoform, the gene model from the *P. sativum* genome (v1a), associated UniProt ID, and corresponding *Arabidopsis thaliana* gene model is listed. Only *P. sativum* gene models assigned to a chromosomal locus were included.

